# Long-term live imaging of the *Drosophila* adult midgut reveals real-time dynamics of cell division, differentiation, and loss

**DOI:** 10.1101/271742

**Authors:** Judy Martin, Erin Nicole Sanders, Paola Moreno-Roman, Shruthi Balachandra, XinXin Du, Leslie Ann Jaramillo Koyama, Lucy Erin O’Brien

## Abstract

Organ renewal is governed by the dynamics of cell division, differentiation, and loss. To study these dynamics in real time, here we present a platform for extended live imaging of the adult *Drosophila* midgut, a premier genetic model for stem cell-based organs. A window cut into a living animal allows the midgut to be imaged while intact and physiologically functioning. This approach prolongs imaging sessions to 12-16 hours and yields movies that document cell and tissue dynamics at vivid spatiotemporal resolution. Applying a pipeline for movie processing and analysis, we uncover new, intriguing cell behaviors: that mitotic stem cells dynamically re-orient, that daughter cells delay fate-determining Notch activation for many hours after birth, and that enterocytes extrude via constriction of a pulsatile cadherin ring. By enabling real-time study of cellular phenomena that were previously inaccessible, our platform opens a new realm for dynamic understanding of midgut organ renewal.

## INTRODUCTION

Stem cell-based organs rely upon the coordinated control of cell division, differentiation, and loss to maintain tissue homeostasis. Studies of the *Drosophila* adult midgut (Fig. 1A) have elucidated conserved processes and pathways that control these events during healthy turnover and cause their dysfunction during aging and in cancer. These contributions, which include mechanisms of multipotency and asymmetric-symmetric fates, endocrine and immune regulation, and injury and stress responses, span the range of adult stem cell biology (Biteau et al., 2008; Buchon et al., 2009; Deng et al., 2015; Guo and Ohlstein, 2015; Hudry et al., 2016; Jiang et al., 2009; O’Brien et al., 2011; Ohlstein and Spradling, 2007; Siudeja et al., 2015).

**Fig. 1.**
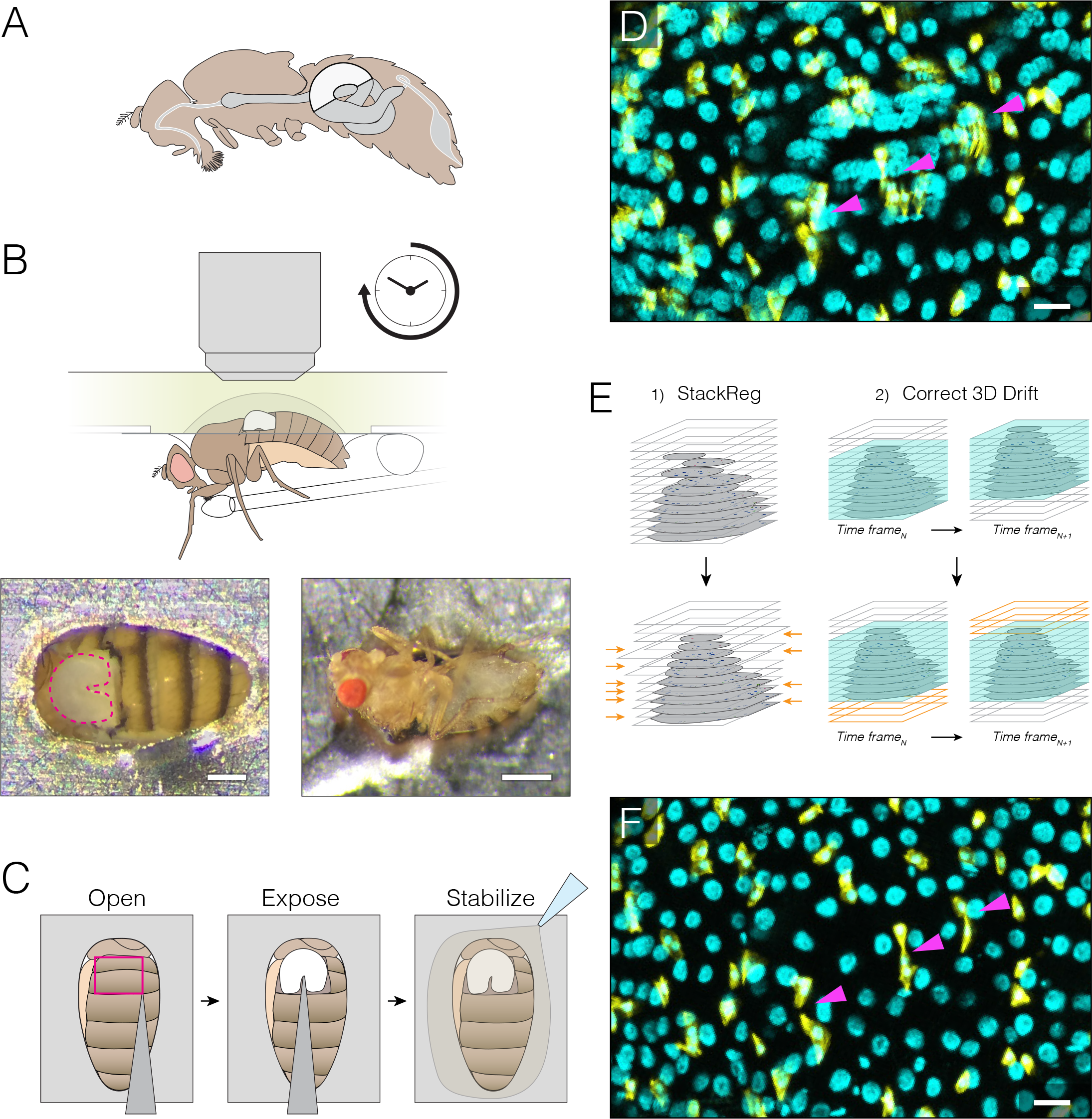
Extended imaging of the midgut in live *Drosophila* adults. **(A)** Adult female midgut *in situ*, sagittal view. White highlight indicates region of the midgut that will be exposed for imaging. **(B-C)** The midgut is accessed through a small cuticular window in the back of a live animal. **B** ***top***, Schematic of imaging apparatus. The animal is affixed to a modified petri dish ‘mount’. The chamber of the mount contains media. The underside of the mount supports a feeder tube. See Fig. S1A-B and S2. **B** ***bottom***, Dorsal (left) and ventral (right) views of an animal in the mount. In the left panel, the exposed midgut is outlined by the magenta dotted line. Scale bars: 0.25 mm (left), 0.5 mm (right). See Video 3. **C**, Steps to prepare the midgut for imaging. See Video 1 tutorial. **(D-F)** Registration macros are applied post-acquisition to correct blurring from tissue movements. **D**, Before registration, blurring and duplications (arrowheads) are evident. Panel is a raw z-series projection of one movie time point. **E**, During registration, two ImageJ plugins are applied in series. (1) StackReg corrects for tissue movement during z-stack acquisition at a single time point. (2) Correct 3D Drift corrects for global volume movements over multiple time points. **F**, After registration, blurring and duplications are negligible. Cyan, all nuclei (*ubi-his2av::mRFP*); yellow, stem cells and enteroblasts (*esg>LifeactGFP*). Scale bars, 20 μm. See Videos 3, 4.

However, investigation of midgut cell dynamics has been constrained by the lack of a viable platform for extended live imaging. At present, fixed midguts provide static snapshots of cells and tissues but do not allow dynamic behaviors to be observed over time. Meanwhile, cultured midguts have been imaged *ex vivo* for 60-90 minutes—a time window long enough for faster events such as calcium oscillations, cell divisions, and acute toxicity responses (Antonello et al., 2015; Deng et al., 2015; Lee et al., 2016; Montagne and Gonzalez-Gaitan, 2014; Scopelliti et al., 2014) but too short for slower, yet also crucial, cell behaviors such as differentiation and apoptosis. Indeed, the power of extended live imaging is demonstrated by studies of numerous other stem cell-based organs, including *Drosophila* ovary and testis (Fichelson et al., 2009; Lenhart and DiNardo, 2015; Morris and Spradling, 2011) and mouse epidermis, testis, muscle, and intestine (Bruens et al., 2017; Gurevich et al., 2016; Hara et al., 2014; Ritsma et al., 2014; Rompolas et al., 2012; Rompolas 2016; Webster et al., 2016). For the midgut, long-term live imaging would synergize with the organ’s existing genetic tractability and well-characterized cell lineages to open exciting investigative possibilities.

To enable such studies, here we present a simple platform that substantially extends imaging times by keeping the midgut within a living animal. The live animal is secured in a petri dish, and the midgut is visualized through a window cut into the dorsal cuticle. The organ’s structural integrity stays largely intact, which allows routine acquisition of movies ~12-16 hours long. Furthermore, digestive function is preserved; the animals ingest food, undergo peristalsis, and defecate even while being imaged. As a result, these long-term movies vividly capture midgut cell dynamics in a near-native physiological context.

To mine the data in these movies, we also present a systematic approach for image processing and segmentation and for spatiotemporal analysis of single cells and whole populations. These proof-of-principle analyses both corroborate prior, fixed-gut observations and reveal intriguing, dynamic behaviors that relate to cell division, differentiation, and loss. (1) For division, we find that mitotic stem cells frequently re-orient—sometimes repeatedly—but can be ‘anchored’ in place by two immature enteroblast cells. (2) For differentiation, we document live activation of a Notch reporter that reveals the transition from a stem-like to a terminal cell state, and we find that, contrary to expectation, real-time activation does not correlate with contact between Notch-and Delta-expressing siblings. (3) For cell loss, we perform morphometric analysis of enterocyte cell extrusion over time and document that two subcellular features of extrusion, nuclear expulsion and constriction of a pulsatile, myosin-rich ring occur with distinct kinetics. Altogether, these analyses demonstrate the potential of long-term live imaging to generate new insights into the dynamics of midgut stem and differentiated cells.

Our study opens the door to live examination of midgut cell dynamics in a near-native context over multi-hour timescales. By allowing real-time observation of cellular events that were previously inaccessible, this platform holds promise to advance understanding of the fundamental cell behaviors that underlie organ renewal.

## RESULTS and DISCUSSION

### An apparatus for midgut imaging within live *Drosophila* adults

We designed a ‘fly mount’ to image the midgut within live *Drosophila* adults (Fig. 1B). Based on a similar design for the brain (Seelig et al., 2010), the midgut mount is assembled from inexpensive, common materials and can be configured for upright, inverted, or light sheet microscopes (Fig. S1A-D). A live animal is stabilized by affixing its abdomen in a ‘mount’ made from a petri dish (up-right or inverted microscopes) or a syringe barrel (light sheet microscopes) (Figs. S1A-D, S2). The midgut is exposed for imaging through a window cut in the dorsal cuticle (Fig. 1B-C). Windows cut in the pliable ventral cuticle were unsuitable for long-term viability. Steps to assemble the fly mount and prepare the midgut are illustrated in a narrated tutorial (Video 1).

Three design features prolong animal viability. First, animals are given liquid nutrients through a feeder tube (Fig. 1B, S1A). Second, the exposed organs are stabilized by a small bed of agarose and bathed in media (Fig. 1C). Third, animals are kept hydrated by a humidity box (Fig. S1B). Even during imaging, animals continue to ingest food, undergo peristalsis, and defecate, which suggests that midguts remain in a state that approaches native physiology.

A crucial element of this setup is the use of a 20x, high-NA dipping objective, which captures z-stacks that are both wide field (100-300 cells) and high resolution (~1 μm) (Video 2). Time intervals between z-stacks ranged from 5-15 min. At room temperature, 72% of animals were alive and responsive after overnight imaging, with typical sessions lasting 12-16 hours (N=18 animals; median imaging duration, 14.6 h) (Video 3). At 29 °C, however, animal viability was substantially shorter, which made temperature-controlled gene expression using GAL80^ts^ impracticable. Progesterone-induced GeneSwitch drivers (Mathur et al., 2010) may provide an alternative.

To minimize interference with native digestion, we used no anesthetics. As a consequence, movies frequently exhibited tissue movements due to both involuntary contractions of the gut tube and voluntary motions of the animal. These movements often resulted in blurred movies that were not amenable to analysis (Fig. 1D, Video 4). However, movement-associated blurring could often be corrected post-acquisition by the sequential application of two ImageJ macros, StackReg and Correct 3D Drift (Fig. 1E) (Arganda-Carreras et al., 2006; Parslow et al., 2014), rendering these movies suitable for single-cell analysis (Fig. 1F, Videos 2, 4).

### A systematic approach for comprehensive spatiotemporal tracking of single cells

Study of dynamic cellular events requires that individual cells be identified, tracked, and analyzed in space and over time. To facilitate these analyses, we generated a ‘fate sensor’ line with fluorescent, nuclear-localized markers for live identification of the midgut’s four major cell types (*esgGAL4, UAShis::CFP*, *GBE-Su(H)-GFP:nls*; *ubi-his::RFP*) (Fig. 2A, B; Video 5). These cell types are: (1) Stem cells, which are marked by CFP and RFP. Stem cells are responsible for virtually all cell divisions. (2) Enteroblasts, which are marked by CFP, GFP and RFP. Enteroblasts are post-mitotic, Notch-activated daughters of stem cells that differentiate into enterocytes. (3) Enterocytes, which are marked by RFP and have polyploid nuclei. Enterocytes are terminally differentiated cells that absorb nutrients and that form the bulk of the midgut epithelium. (4) Enteroendocrine cells, which are marked by RFP and have small, diploid nuclei. Enteroendocrine cells are terminally differentiated cells that secrete enteric hormones.

**Fig. 2.**
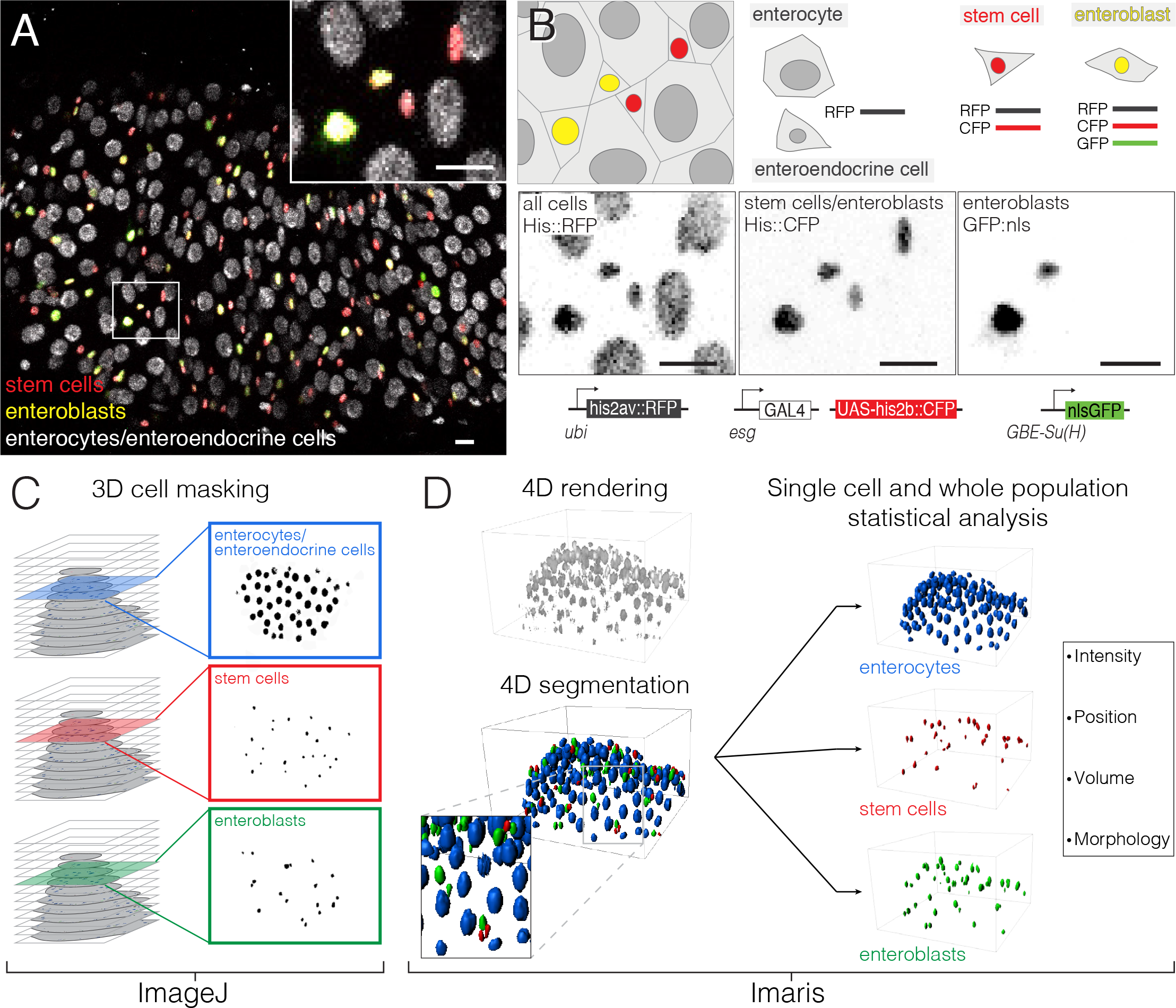
Comprehensive, fate-specific tracking and analysis of individual cells. **(A-B)** ‘Fate sensor’ midguts enable live identification of cell types. **A**, Stack projection of a single time point from a 10-hour movie (Video 5). Nuclei are distinguishable for four midgut cell types: stem cells (red pseudocolor), enteroblasts (yellow-green pseudocolor), enterocytes (gray, polyploid), and enteroendocrine cells (gray, diploid). Inset shows the zoom region depicted in B. **B**, Genetic design of fate sensor line (*esg>his2b::CFP*, *GBE-Su(H)-GFP:nls*; *ubi-his2av::mRFP*). Cell types are distinguished by combinatorial expression of three fluorescent, nuclear-localized markers: enterocytes/enteroendocrine cells (His2ab::mRFP only), stem cells (His2ab::mRFP, His2b::CFP), and enteroblasts (His2ab::mRFP, His2b::CFP, GFP:nls). All scale bars, 10 μm. **(C-D)** Workflow to identify, track, and analyze cells in volumetric movies. **C**, From raw, multi-channel z-stacks, nuclei are digitally separated into stem cell, enteroblast, and enterocyte/entero-endocrine populations using channel masks in ImageJ. **D**, The three population sets are rendered in 4D in Imaris. Segmentation is performed on each population to identify individual nuclei. Entero-endocrine nuclei are separated from enterocyte nuclei by a size filter. Positions of individual nuclei are correlated between time points to track single cells over time.

To analyze these multichannel, volumetric movies, we developed a semi-automated workflow that uses ImageJ and Bitplane Imaris to digitally separate marked populations, identify all individual cells in each population, and track these cells for the duration of the movie (Fig. 2C-D). This comprehensive single-cell tracking enables features such as fluorescence intensity, spatial position, and nuclear size to be measured for each individual cell. Multiplying the 100-300 cells in a movie over the hundreds of time points captured in a 12-16 hour imaging session, we collect tens of thousands of real-time measurements. Unlike prior approaches, which used manual identification and tracking of selected cells, our comprehensive approach provides both single-cell and population-level data in an unbiased manner.

To demonstrate the utility of this imaging platform and workflow, we performed proof-of-principle analyses for three core behaviors of midgut renewal: enterocyte extrusion and loss (Fig. 3A-F), stem cell division (Figs. 3G-H, 4), and enteroblast differentiation (Figs. 5-6).

**Fig. 3.**
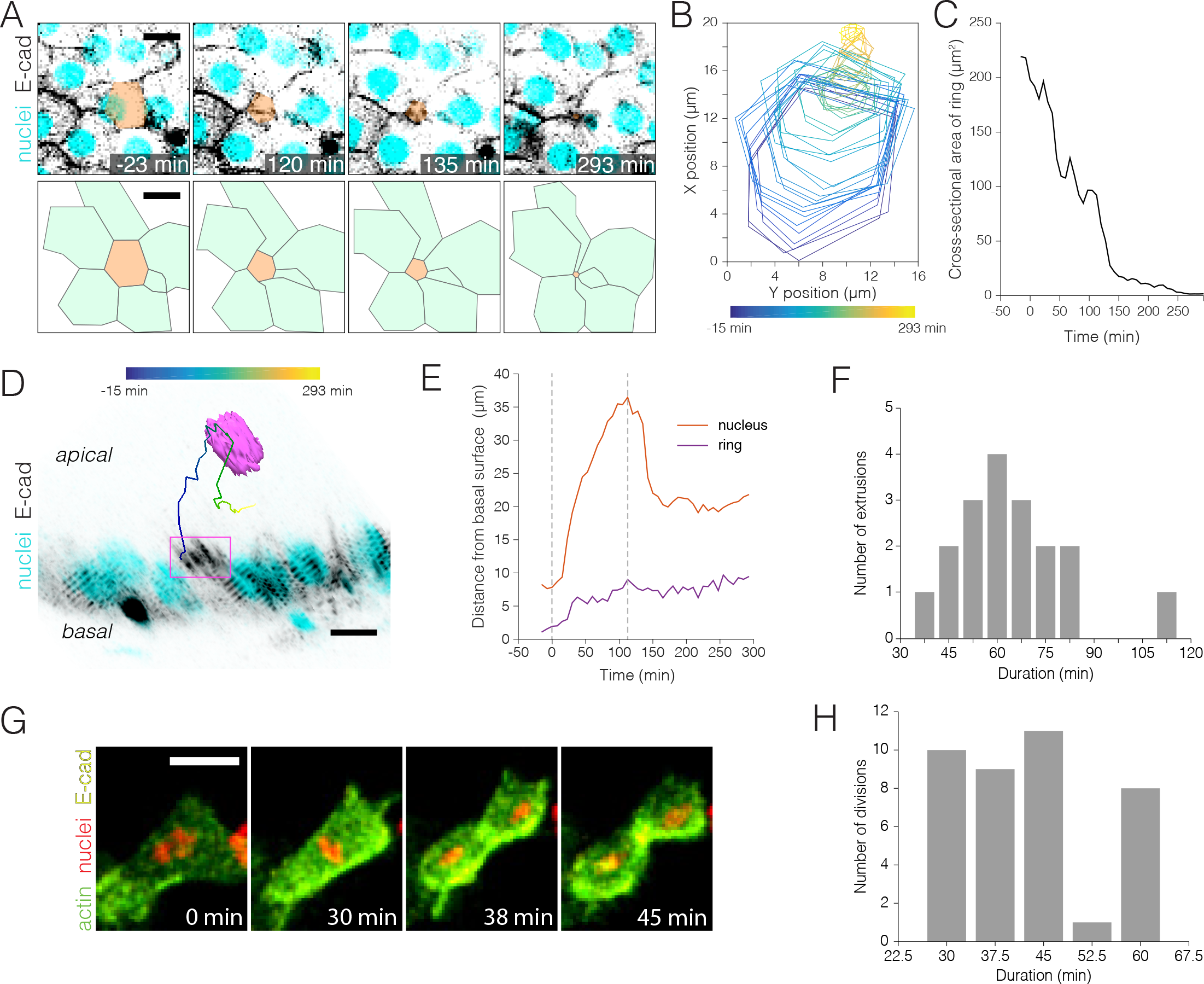
Real-time kinetics of enterocyte extrusion and stem cell mitosis. **(A-E)** Proof-of-principle analysis of a single-enterocyte extrusion. **A**, Time-lapse sequence (top) and schematic (bottom) of an extrusion event. The basal region of the extruding cell (tan pseudocolor) is outlined by a six-sided ‘ring’ of E-cadherin::YFP (inverted gray, *ubi-DE-cadherin::YFP*). Over time, the ring closes to a point, and the six neighbor cells (green in schematic) draw into a rosette. Top panels are stack projections. Cyan (*ubi-his2av::mRFP*) labels all nuclei. See Video 6. **B**, Spatial ‘foot-print’ of the E-cadherin::YFP ring in the epithelial plane over time (violet-yellow color scale). The ring remains six-sided throughout closure. **C**, Kinetics of ring closure are pulsatile. As the cross-sectional area of the ring shrinks, long phases of constriction are interrupted by shorter phases of relaxation. **D**, The extruding nucleus (magenta) ejects into the lumen then recoils. Orthoview of the same extrusion as A. Multicolored line shows path of nucleus over time (violet-yellow color scale). Magenta box denotes the E-cadherin::YFP ring, which is visible in this time point (t=135 min) as a density of YFP at the apical surface. Inverted gray, E-cadherin::YFP (*ubi-DE-cadherin::YFP*); cyan, all nuclei (*ubi-his2av::mRFP*). See Video 7. **E**, The nucleus and ring move apically with distinct ki-netics. Displacement from the basal surface over time is shown for the ring (purple) and nucleus (red). The ring advances incrementally via small, apical-and-basal movements. The nucleus shoots rapidly into the lumen, then recoils. **F**, Durations of nuclear extrusion for 18 single enterocytes from 6 movies. Nuclear extrusions range from 37-112 min with a mean of 64 ± 18 min (SD). **(G-H)** Kinetics of stem cell mitoses. **G**, Time-lapse sequence of a mitotic event. Green, actin (*esg>LifeactGFP*); yellow, E-cadherin (*ubi-DE-cadherin::YFP*); red, nuclei (*ubi-his2av::mRFP*). Panels are partial stack projections of the basal epithelium. See Video 8. **H**, Durations of mitosis for 39 cell division from 11 movies. Mitoses range from 30-60 min with a mean of 43 ± 11 min (SD). All scale bars, 10 μm.

### Enterocyte extrusion: Spatiotemporal dynamics of ring closure, ring travel, and nuclear travel

Enterocytes in the *Drosophila* midgut, like enterocytes in the mammalian intestine, are lost through apical extrusion (Buchon et al., 2010; Eisenhoffer et al., 2012; Harding and Morris, 1977; Madara, 1990; O’Brien et al., 2011). During extrusion, a cell is ejected out of the epithelium and into the lumen through the concerted contractions of its neighbors (Eisenhoffer and Rosenblatt, 2012). Because this process is seamless, extrusion allows apoptotic cells to be eliminated while still preserving the epithelial barrier (Gudipaty and Rosenblatt, 2017). Since apoptotic enterocytes secrete stem cell-activating mitogens (Liang et al., 2017), understanding when and how apoptotic enterocytes are extruded is important for understanding the dynamics of midgut turnover.

Extrusion has been challenging to study in fixed tissues because extruded cells leave no trace in the epithelium. Fixed sections can occasionally catch cells ‘in the act’ of extruding, but they do not reveal the dynamics of these transient events.

Our imaging platform provided the opportunity to study *in vivo* extrusions live. We observed that enterocytes extrude either as single cells (18 of 34 extrusions; Fig. 3A, F; Videos 6, 7) or as groups of 2-5 cells (16 of 34 extrusions). We also observed one instance in which diploid cell extruded. All extrusions were apical.

Single-enterocyte extrusions had two prominent dynamic features: (1) Closure and apical movement of cortical, contracile ‘ring’ (Fig. 3A-C, E, Video 6). (2) Luminal ejection then recoil of the nucleus (Fig. 3D-F, Video 7). As a case study to highlight the spatiotemporal resolution of our data, we performed morphometric analysis of the extrusion in Fig. 3A.

The basal region of this extruding enterocyte was outlined by a six-sided ‘ring’ of E-cadherin::YFP (Fig. 3A). The ring’s changing size and position sheds light on the kinetics of basal detachment and delamination. Over a 5 h-period, the ring closed to a point and eventually vanished (Fig. 3A-C; Video 6), suggesting that the basal region of the extruding cell was shrinking and ultimately expelled. Throughout much of extrusion, the intensity of E-cadherin::YFP fluctuated (Video 6, t=-150-134 min), perhaps reflecting phases of constriction and relaxation (see below). However, as the ring drew tightly to a point, the E-cadherin signal became consistently bright (Video 6, t=135-293 min). As the ring closed, its six surrounding cells formed a rosette with the extruding cell at its center.

Measuring the cross-sectional area of the ring over time, we observed that ring closure was pulsatile. Longer phases of constriction, lasting 1-2 h, were interrupted by shorter phases of relaxation, lasting just 20-30 min (Fig. 3D). This pulsatile pattern, which has not been observed in cultured epithelial cells (Kuipers et al., 2014), is reminiscent of the acto-myosin ‘ratcheting’ that co-ordinates junctional remodeling of developing epithelia (Coravos et al., 2017).

Both the ring and the extruding cell’s nucleus moved from the basal epithelial surface toward the apical lumen. However, their kinetics of travel were distinct (Fig. 3E). The ring advanced slowly, with stuttering, apical-and-basal movements that produced net apical progress over 5 h. By contrast, the nucleus shot rapidly out of the epithelium over 15 min (Fig. 3D; Video 7, t=0-15min) and continued to penetrate deeper into the lumen over the next ~1.6 h (Fig. 3D; Video 7, t=15-113 min). After reaching its maximum luminal depth, the nucleus recoiled and came to rest on the apical epithelium (Fig. 3D; Video 7, t=113-293 min). Thus, ring and nuclear travel occur with distinct kinetics, which suggests that these two subcellular events involve different mechanisms.

Altogether, this detailed case study illustrates the high-resolution cellular morphometrics that are possible with our movie data. Qualitative examination of two other single-enterocyte extrusions suggested similar ring and nuclear dynamics, providing a basis for future investigations. By enabling direct comparison of concurrent subcellular events in real time, our platform provides a new, dynamic view of epithelial cell extrusion in a physiologically functioning organ.

### Stem cell division: Mitotic orientation in real time

Tissue homeostasis requires the replacement of extruded cells by new cells. In the adult *Drosophila* midgut, new cells are generated through stem cell divisions; terminal daughters are typically post-mitotic. Although time-lapse imaging has unique potential to reveal division behaviors (Park et al., 2016), the divisions of midgut stem cells have been challenging to capture. To date, live divisions have been reported in only one study, which examined pathogen-stimulated midguts *ex vivo* (Montagne and Gonzalez-Gaitan, 2014).

We surveyed our movies of near-native midguts for physiological divisions. Thirty-nine mitoses were identified in 11 movies with a combined duration of 122 hours. Within individual movies, mitoses were distributed similarly between the first and second halves of a movie. No divisions were identified in 14 additional movies. The average mitosis lasted 43 ± 11 min (Fig. 3G, H), and each mitosis was exhibited by a unique stem cell. From these findings, we calculated a mitotic index of 0.28% (see Methods), which is less than the 1-2% mitotic index obtained from counts of phos-pho-histone H3^+^ stem cells in fixed midguts (Jin et al., 2017; Kolahgar et al., 2015; Montagne and Gonzalez-Gaitan, 2014). Adjustments to media formulations or imaging conditions might help to reduce this difference.

We investigated how mitotic stem cells dynamically orient in 3D space. In general, dividing epithelial cells orient themselves in two frames of reference: horizontal-vertical and longitudinal-circumferential. Horizontal-vertical orientation is defined by the epithelial plane (Fig. 4A) and, in development, serves to determine daughter fates (Cayouette and Raff, 2003; Dong et al., 2012; El-Hashash et al., 2011; Guo and Ohlstein, 2015; Williams et al., 2011). Longitudinal-circumferential orientation is defined by organ shape (Fig. 4F) and determines whether the organ grows longer or wider (Mochizuki et al., 2014; Tang et al., 2011). In developing epithelia, well-understood regulatory mechanisms ensure that divisions place new cells in the right positions for proper organ morpho-genesis. In mature, physiologically functioning epithelia, however, whether or not such mechanisms persist is an important open question.

**Fig. 4.**
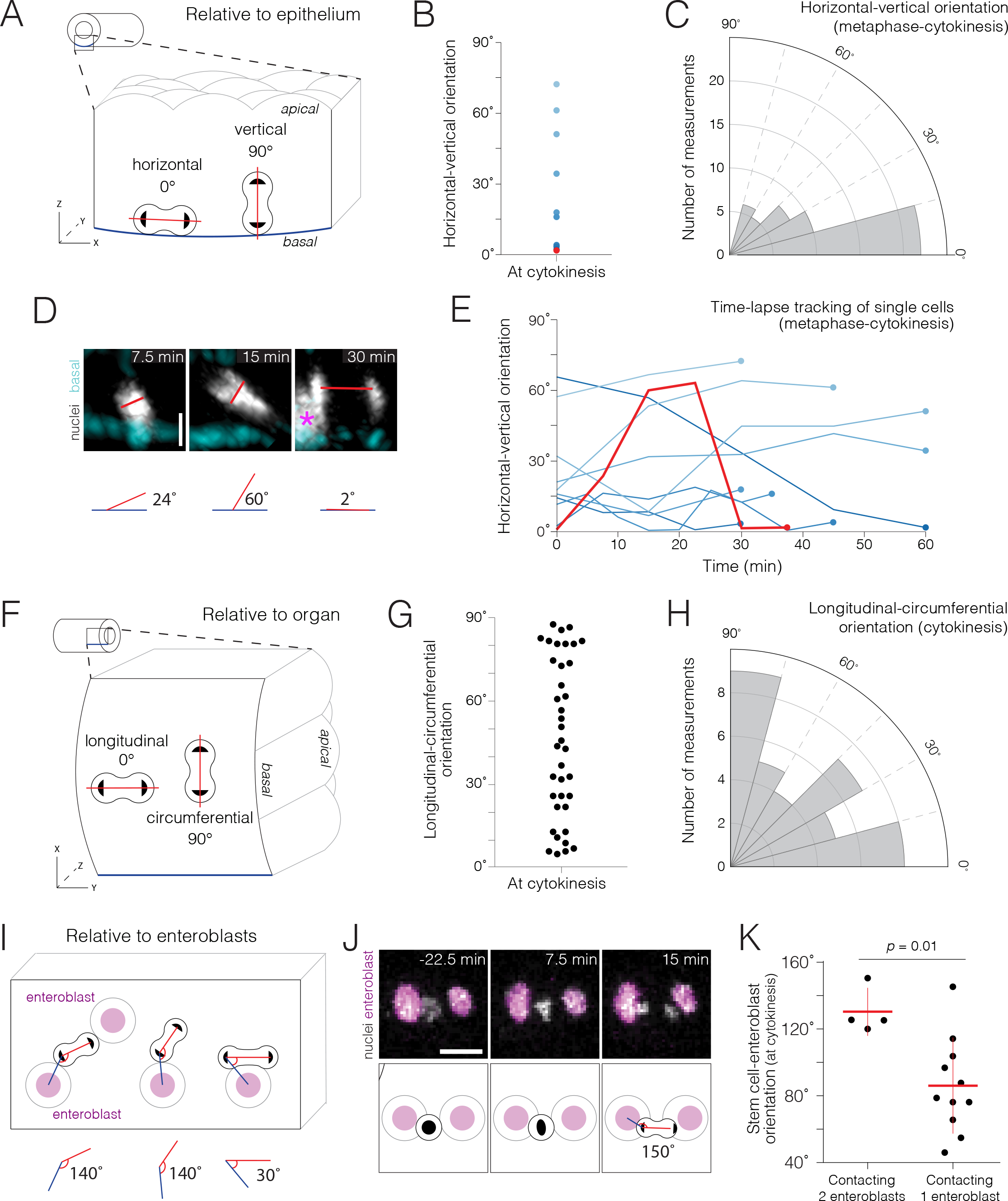
Real-time orientations of stem cell divisions in three reference frames. **(A-E)** Horizontal-vertical orientations are horizontally biased. **A**, Schematic of horizontal (0°) and vertical (90°) orientations. See Fig. S3. **B**, Live orientations of 10 dividing cells specifically at cytokinesis. The distribution is biased toward horizontal (<45°). Red point represents the cell in Panel D. **C**, Live orientations of the same 10 cells throughout mitosis. Each measurement is the orientation of one mitotic cell at one time time point, from metaphase to cytokinesis (n=51 measurements). The distribution is biased toward horizontal (<45°). **D**, Two re-orientations in a single mitosis. Red line shows the orientation of condensed chromatin (gray, *ubi-his2ab::mRFP*) relative to the basal basement membrane (cyan, Concanavalin A-Alexa-647). For clarity in the 7.5 min and 15 min projections, a clipping plane was applied in the gray channel to exclude an enterocyte nucleus; this nucleus is marked by an asterisk at the left edge of the 30 min projection. Scale bar, 5 μm. See Video 9. **E**, Mitotic cells frequently re-orient. Each line shows the horizontal-vertical orientations of a single mitotic cell over time. The 10 cells are the same as in Panels B and C. All lines start at metaphase (t=0 min) and continue to cytokinesis (t=30-60 min). Time intervals were either 5, 7.5, or 15 min. Colors are the same as in Panel B; red line is the cell in Panel D. **(F-H)** Longitudinal-circumferential orientations are unbiased. **F**, Schematic of longitudinal (0°) and circumferential (90°) orientations. **G-H**, Live orientations of 38 dividing cells at cytokinesis. Longitudinal (≤45°) and circumferential (>45°) orientations are near-equal. **(I-K)** Divisions between two flanking enteroblasts align with the enteroblast-enteroblast axis. **I**, Schematic of divisions contacting either two or one enteroblast(s). When two enteroblasts are present, the closer enteroblast is used for measurements (see Methods). **J**, Division between two enteroblasts. Orientation is nearly parallel to the axis between the enteroblast nuclei (magenta, *GBE-Su(H)-GFP:nls*). Gray, stem cell and enteroblast nuclei (*esg>his2b::CFP*). Scale bar, 10μm. See Video 10. **K**, Live orientations of divisions with two or one flanking enteroblast(s). With two enteroblasts (n=4 of 18 divisions), orientations are near-parallel to the enteroblast-enteroblast axis. With one enteroblast (n=11 of 18 divisions), orientations are broadly distributed. Orientations were measured at cytokinesis. Means ± SD are shown. Mann-Whitney test, *p*=0.01.

To shed light on this question, we investigated whether or not the native mitoses of midgut stem cells exhibited bias in either horizontal-vertical or longitudinal-circumferential orientations. While performing these two analyses, we noticed an unexpected, third frame of reference: neighbor enteroblasts (Fig. 4I). Below, we describe real-time mitotic orientation in these three reference frames.

First, we considered the horizontal-vertical reference frame (Fig. 4A-E). At cytokinesis, hori-zontal-vertical orientations ranged broadly, from 1.6°-72°, but were biased toward horizontal; of 10 dividing cells, 4 were <5° and 7 were <45° (Fig. 4B). These findings are consistent with prior analyses of dividing stem cells in fixed midguts (Goulas et al., 2012; Ohlstein and Spradling, 2007).

Do horizontal-vertical orientations stay constant throughout division? Analyzing the same 10 cells from metaphase to telophase, we found that orientations were, as a group, also biased toward horizontal; of 41 measurements, 7 were <5° and 33 were <45° (Fig. 4C). Interestingly, however, this stable, population-level trend belied the dynamic re-orientations of individual cells. Tracking single cells over time, we found that 8 of the 10 cells re-oriented by ≥15° at least once, and 4 re-oriented by ≥30° (Fig. 4D, E). Re-orientations even occurred repeatedly during a single mitosis; 3 cells re-oriented by ≥15° two or three times (Fig. 4D, E). These frequent, sometimes dramatic, re-orientations are detectable uniquely by live imaging and carry implications for how mitotic orientations in fixed midguts are interpreted.

Second, we considered the longitudinal-circumferential reference frame (Fig. 4F-H). Thirty-eight mitotic cells were examined specifically at cytokinesis. In this sample, longitudinal and circumferential orientations occurred with near-equal frequency; 20 cells were ≤45° and 18 cells were >45° (Fig. 4F-H). These data indicate that, unlike horizontal-vertical orientations, longitudinal-circumferential orientations are unbiased.

Finally, we observed that a third, local reference frame was formed when two enteroblasts flanked a stem cell (Fig. 4I-K). In this three-cell arrangement, divisions occurred nearly parallel to the two neighbor enteroblasts (4 of 18 divisions; Fig. 4J, K). By contrast, divisions exhibited a broad range of orientations if only one neighbor enteroblast was present (11 of 18 divisions; Fig. 4K). When trapped between two enteroblasts, daughter cells at cytokinesis hurled into and forcibly collided with the enteroblast nuclei (Fig. 4J; Video 10, t=15-22.5 min). These observations suggest that contacts between stem cells and enteroblasts provide a spatial cue that orients the mitotic spindle.

In summary, our analyses provide the first views of how live stem cells orient their divisions within the midgut’s tubular epithelium. They also reveal mitotic behaviors, such as frequent horizontal-vertical re-orientations, that are indiscernible in fixed samples. Examining dividing stem cells in three reference frames, we found three orientation patterns: (1) Biased horizontal orientations. In future work, a crucial question will be whether, and if so how, horizontal orientations promote symmetric daughter fates (Goulas et al., 2012; Guo and Ohlstein, 2015; Kohlmaier et al., 2015; Montagne and Gonzalez-Gaitan, 2014; Sallé et al., 2017). (2) Unbiased longitudinal-circumferential orientations. This balanced distribution may help maintain constant organ shape over time. (3) Local orientation by two enteroblasts. This unanticipated finding raises the possibility that stem cell-enteroblast adherens junctions, which are unusually pronounced (Ohlstein and Spradling, 2006), may orient the divisions of midgut stem cells similar to the adherens junctions of Drosophila male germline stem cells, sensory organ precursors, and neuroblasts (Inaba et al., 2010; Le Borgne et al., 2002; Lu et al., 2001). These findings provide a basis for future work to directly investigate the mechanisms that orient midgut stem cells and to probe the relationship between division orientation and daughter fates.

### A quantitative threshold of Notch activation distinguishes stem cells and enteroblasts

Along with cell division and loss, cell differentiation is the third core behavior of tissue renewal. In the *Drosophila* adult midgut, differentiation in the enteroblast-enterocyte lineage is controlled by Delta-Notch. Delta ligand, which is predominantly on stem cells, activates Notch receptor on stem (or stem-like) cells. At low levels, Notch activity is compatible with stemness. However at higher levels, it triggers enteroblast differentiation (Bardin et al., 2010; Biteau and Jasper, 2014; Kohlmaier et al., 2015; Micchelli and Perrimon, 2006; Ohlstein and Spradling, 2006; 2007; Perdigoto et al., 2011; Zeng and Hou, 2015).

Movies of ‘fate sensor’ midguts provide a means to measure Notch activity live (Fig. 2). In these guts, the *GBE-Su(H)-GFP:nls* reporter provides a sensitive readout of Notch transcriptional activity (de Navascués et al., 2012; Furriols and Bray, 2001; Guisoni et al., 2017; Housden et al., 2014), while *ubi-his2av::mRFP* provides a stable reference signal. We used these two markers to establish a quantitative metric of Notch activation that enables comparison of the same cell at different times, or of different cells within the same movie or across different movies. First, to account for differences between movies, we normalized the values of GFP and RFP intensities within a given movie to a 0-to-1 scale. Second, to account for tissue depth and other artifacts within a single movie, we used these normalized GFP and RFP intensities to calculate the ratio of GFP:RFP for each *esg>his2ab::CFP* progenitor cell at each time point. This two-part calculation of real-time GFP:RFP enables Notch activity to be compared over time and between cells.

We asked whether real-time GFP:RFPs are consistent with conventional indicators of enteroblast differentiation. GFP:RFPs, which ranged from 0.0-1.8, generally fit with subjective evaluations of GFP intensities (Figs. 5A, B). In addition, population-level distributions of GFP:RFP were similar at different imaging depths, over time within a single movie, and across different movies. Furthermore, *esg*^+^ cells with large nuclei (≥200 μm^3^) often exhibited high GFP:RFPs, while cells with low GFP:RFP typically had small nuclei (Fig. 5B). This association of high GFP:RFPs with large but not small enteroblast nuclei fits with prior observations that endoreplication is specific to late enteroblasts (Jiang et al., 2009; Kohlmaier et al., 2015; Lucchetta and Ohlstein, 2017; Perdigoto et al., 2011; Xiang et al., 2017; Zhai et al., 2017). Altogether, these findings support the use of real-time GFP:RFPs as a metric of Notch reporter activation.

**Fig. 5.**
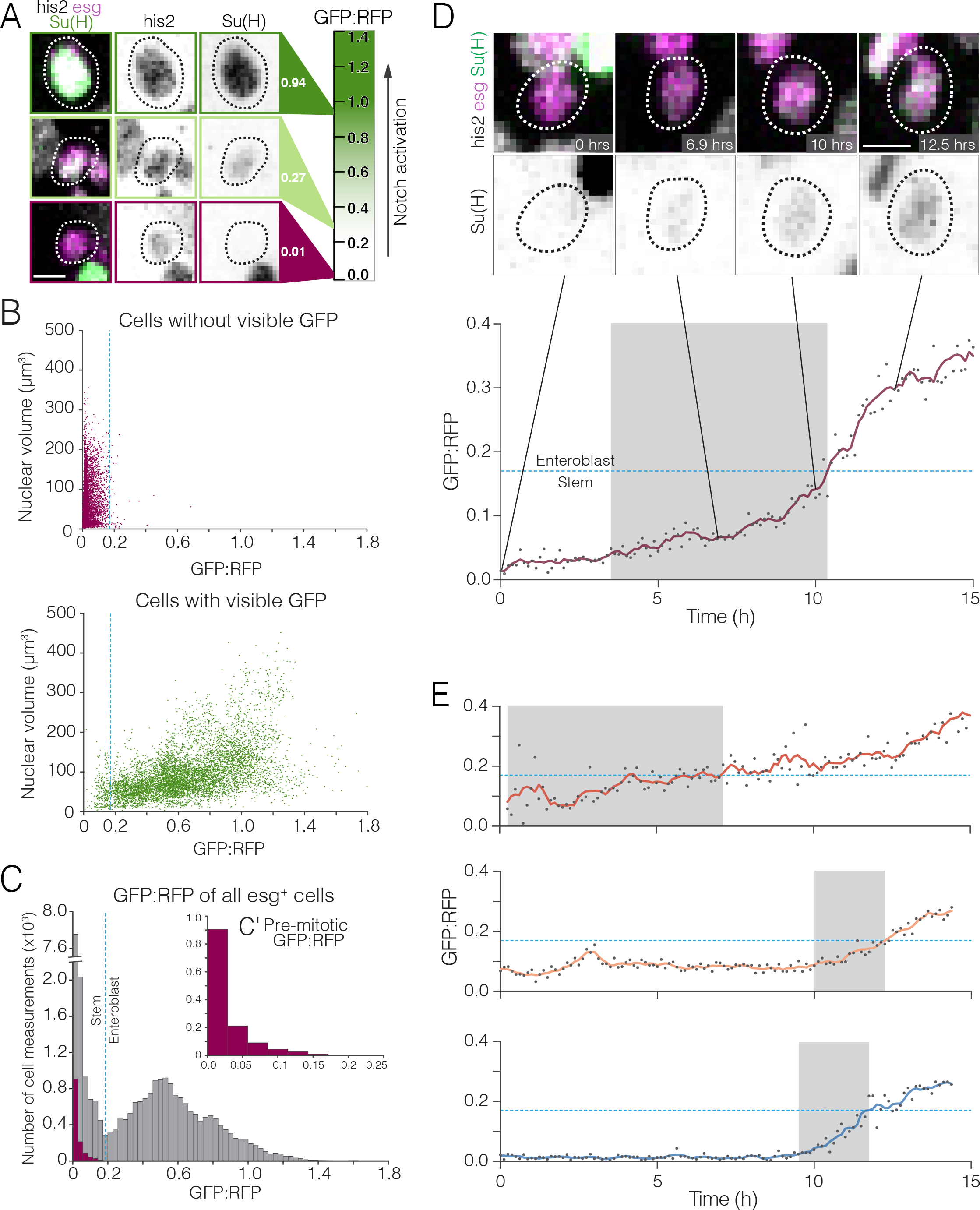
Whole-population and single-cell analyses of real-time Notch activation. **(A-C)** A threshold level of Notch activation distinguishes stem cells and enteroblasts. **A**, Single-cell measurements of the Notch reporter *GBE-Su(H)-GFP:nls* from live movies. Cells additionally co-express *esg>his2ab::CFP* (magenta) and *ubi-his2ab::mRFP* (gray). *GBE-Su(H)-GFP:nls* activation is quantified as GFP:RFP (see Methods). For the indicated cells, GFP:RFP=0.94, 0.27, and 0.01. **B**, GFP:RFP values correlate with visible GFP and nuclear volume. Progenitor (*esg*^+^) cells were scored by eye as either GFP-negative (top) or -positive (bottom). In cells without visible GFP, nearly all GFP:RFP values cluster between 0.0 and 0.2, and most nuclear volumes are small (<200 μm_3_). In cells with visible GFP, most GFP:RFP values are spread between 0.1 and 1.4, and large nuclear volumes (≥200 μm^3^), indicative of late enteroblasts, are associated with high GFP:RFPs. Blue dotted line shows the 0.17 enteroblast threshold from Panel C. C, GFP:RFP values quantitatively distinguish stem cells and enteroblasts. Gray bars show real-time GFP:RFPs for all *esg*^+^ cells in two movies (29,102 GFP:RFPs from 251 cells). Two peaks (GFP:RFP=0.015, 0.528) are separated by a local minimum (blue dotted line; GFP:RFP=0.17). Purple bars (C′ inset) show real-time GFP:RFPs for ‘benchmark’ stem cells prior to an observed mitosis (1,294 GFP:RFPs from 18 premitotic cells). The benchmark stem cell distribution matches the left peak of the *esg*^+^ cells, and 99.6% of ‘benchmark’ GFP:RFPs are less than 0.17. Data in B and C are the aggregate of 2 movies. **(D-E)** Stem-like cells transition to enteroblasts over multiple hours. **D**, Real-time activation of *GBE-Su(H)-GFP:nls* reveals a transition from a stem-like to an enteroblast state. During a transition period lasting 6.9 h (gray background), GFP:RFP increases from a baseline of ~0.049 at t=3.5 h to the enteroblast threshold of 0.17 (blue dotted line) at t=10.4 h. After the transition, GFP:RFP continues to increase and reaches 0.364 at t=15.0 h. *GBE-Su(H)-GFP:nls* shown in green (top) and inverted gray (bottom); *esg>his2ab::CFP*, magenta; *ubi-his2ab::mRFP*, gray. See Video 11. E, Kinetics of three additional enteroblast transitions. Initial baseline GFP:RFPs are <0.17. GFP:RFPs increase from baseline to 0.17 during transition periods lasting from 2.3-6.9 h (gray backgrounds: t=0.3-7.1 h (top), 10.0-12.4 h (middle), 9.5-11.8 h (bottom)). Initial and final GFP:RFPs are as follows: 0.058, 0.426 (top); 0.069, 0.281 (middle); 0.022, 0.257 (bottom). All cells in Panels D and E were born before imaging started. Genotype in all panels: *esgGal4*, *UAS-his2b::CFP*, *Su(H)GBE-GFP:nls*; *ubi-his2av::mRFP*. All scale bars are 5 μm.

Having validated real-time GFP:RFPs, we used them to investigate the different levels of Notch activation that are associated with stem and enteroblast identity. The existence of these different levels was originally identified via genetic modulation of Notch signaling (Perdigoto et al., 2011), but actual levels of Notch signaling had not been evaluated quantitatively. We wondered whether GFP:RFP values could distinguish stem cells and enteroblasts in real time.

Examining GFP:RFPs for all *esg*^+^ progenitor cells in two fate sensor movies (29,102 values; 251 cells), we found that their distribution is—suggestively—bimodal. A local minimum at GFP:RFP=0.17 separates a sharp left peak (GFP:RFP=0.015) and a broad right peak (GFP:RFP=0.528) (Fig. 5C). An appealing interpretation of this bimodality is that the left peak represents stem cells and the right peak represents enteroblasts.

To directly test this interpretation, we cross-correlated real-time Notch activity with mitotic behavior. Mitosis is near-exclusive to stem cells, so cells that exhibited mitosis are identifiable as stem cells irrespective of their GFP:RFPs. The GFP:RFPs of these cells *in the time points prior to their observed mitoses* thus represent a ‘benchmark’ collection of GFP:RFPs from known stem cells (1,294 GFP:RFPs; 18 cells).

The benchmark stem cell GFP:RFPs were compared to all progenitor GFP:RFPs. If the left peak of the bimodal *esg*^+^ distribution represents stem cells, then its GFP:RFP profile should resemble the profile of the ‘benchmark’ stem cells. Indeed, the two profiles nearly matched. Furthermore, 99.61% of GFP:RFPs for benchmark stem cells were less than the 0.17 threshold (Fig. 5C′). This correspondence implies that stem cells populate the left peak and that enteroblasts populate the right peak. Supporting this interpretation, the number of data points in the left and right peaks have a proportion of 4:3, which is similar to previously reported proportions of stem cells to enteroblasts in fixed tissues (Guisoni et al., 2017; O’Brien et al., 2011). Based on these findings, we conclude that GFP:RFP=0.17 represents a threshold of Notch activation that functionally distinguishes stem cells and enteroblasts in fate sensor movies.

### Real-time kinetics of enteroblast transitions

Having identified this stem-enteroblast threshold, we applied it to investigate how cells transition from stem-like to enteroblast states. A fundamental aspect of fate transitions is the time over which they occur. Fast transitions would allow cells to respond nimbly to acute challenges, whereas slow transitions would allow cells to receive and integrate a large number of fate-influencing signals. In this manner, the kinetics of fate transitions can define how an organ responds to changing external environments.

Midgut fate transitions have not been measured directly to date. For enteroblasts, an upper limit of two days can be inferred from observations that enteroblasts are present in stem cell clones two days post-induction (de Navascués et al., 2012; Ohlstein and Spradling, 2007). However, in developing tissues such as the vertebrate neural tube and the *Drosophila* peripheral nervous system and wing imaginal disc, activation of Notch target genes occurs over minutes-to-hours (Corson et al., 2017; Couturier et al., 2012; Housden et al., 2013; Vilas-Boas et al., 2011). Notch kinetics in these other tissues raise the possibility that Notch-mediated enteroblast transitions in the midgut could be considerably faster than two days.

To examine these kinetics directly, we measured the rate of *GBE-Su(H)-GFP:nls* activation in movies of fate sensor midguts. Cells were scored as undergoing an enteroblast transition if their GFP:RFPs persistently increased from an initial baseline of GFP:RFP <0.17 to a value greater than the 0.17 enteroblast threshold. From 95 cells with initial GFP:RFP<0.17 in two movies, 5 such transitions were identified. We focused initially on 4 of these, each of which occurred in a cell that was born before imaging started (Fig. 5D-E, Video 11). In each of these 4 instances, GFP:RFP started from a different baseline and increased at a different rate. Nonetheless in all cases, the actual transition occurred over multiple hours (2.4-6.9 h; gray background shading in Fig. 5D-E). Interestingly, we also observed *esg*^+^ cells in which GFP:RFP fell from above to below 0.17 over several hours. These events might suggest that some progenitors activate Notch to enteroblast levels before reverting to a stem-like state. Altogether, our real-time data reveal that stem-like cells activate a Notch reporter and differentiate into enteroblasts on a time scale of hours—much faster than the 2-day upper limit implied by clones, and much slower than the minutes observed in some other tissues. These analyses demonstrate the ability of our imaging platform to provide new insights on the dynamic nature of stem and enteroblast identity.

### Sibling contacts do not instigate real-time Notch activation

In order to activate Notch, a potential enteroblast must have physical contact with a Delta-expressing stem cell. In principle, newborn sibling cells are ideal partners to engage in Notch-Delta interactions with each other (Guisoni et al., 2017). Newborn cells express both Notch and Delta (Bardin et al., 2010; Ohlstein and Spradling, 2007), and cytokinesis leaves sibling cells juxtaposed. For Notch to become activated, siblings must remain in contact long enough to overcome the time delays that are inherent to Delta-Notch lateral inhibition (Barad et al., 2010; Du et al., 2017; Guisoni et al., 2017). However, fixed-gut studies of twin spot clones imply that after cytokinesis, some sibling pairs become separated (O’Brien et al., 2011). If sibling contacts can be transient, then the relationship between contact dynamics and Notch activation kinetics becomes a crucial variable in enteroblast specification.

To investigate contact dynamics, we sought to visualize them directly by incorporating a membrane-localized YFP into our fate sensor line, which already contained nuclear-localized CFP, GFP, and RFP (Fig. 2A-B). However, we were unable to parse the YFP signal without sacrificing sensitivity in the critical GFP channel for the Notch reporter. As an alternative approach, we evaluated whether contact between siblings could be inferred from the distance separating their nuclei. To directly compare cell-cell contact and inter-nuclear distance in the same progenitor pairs, we used movies of midguts in which progenitor cell boundaries were visualized by LifeactGFP and nuclei were visualized by His2av::RFP (Fig. S4). This analysis revealed two strong correlations: progenitors with inter-nuclear distance <6.0 μm were nearly always in contact, and progenitors with inter-nuclear distance >15.5 μm were nearly always separated. Inter-nuclear distances between 6.0-15.5 μm did not correlate with either contact or separation. Based on these findings, we designated three classifications: inferred contact (inter-nuclear distance <6.0 μm), indeterminate (inter-nuclear distance ≥6.0 and ≤15.5 μm), and inferred separation (inter-nuclear distance >15.5 μm).

We used these classifications to examine the contact dynamics of sibling pairs in movies of fate sensor midguts. Analyzing 18 sibling pairs with known birth times, we found they exhibited a broad diversity of contact behaviors (Fig. 6A). At one extreme were high-contact pairs, which generally stayed in place after cytokinesis (pairs A-C, Fig. 6A). At the other extreme were low-contact pairs, which separated within minutes after cytokinesis (pairs N-R, Fig. 6A). Eleven of 18 pairs separated for one or more hours; two pairs separated and contacted repeatedly; and six pairs appeared to separate permanently. Notably, these dynamic, variable contact behaviors could not have been deduced from static images. Our observations that siblings lose contact with surprising frequency suggest that a substantial proportion of true sibling pairs may be missed by conventional fixed-gut assays that consider only *esg*^+^ cell pairs as siblings.

**Fig. 6.**
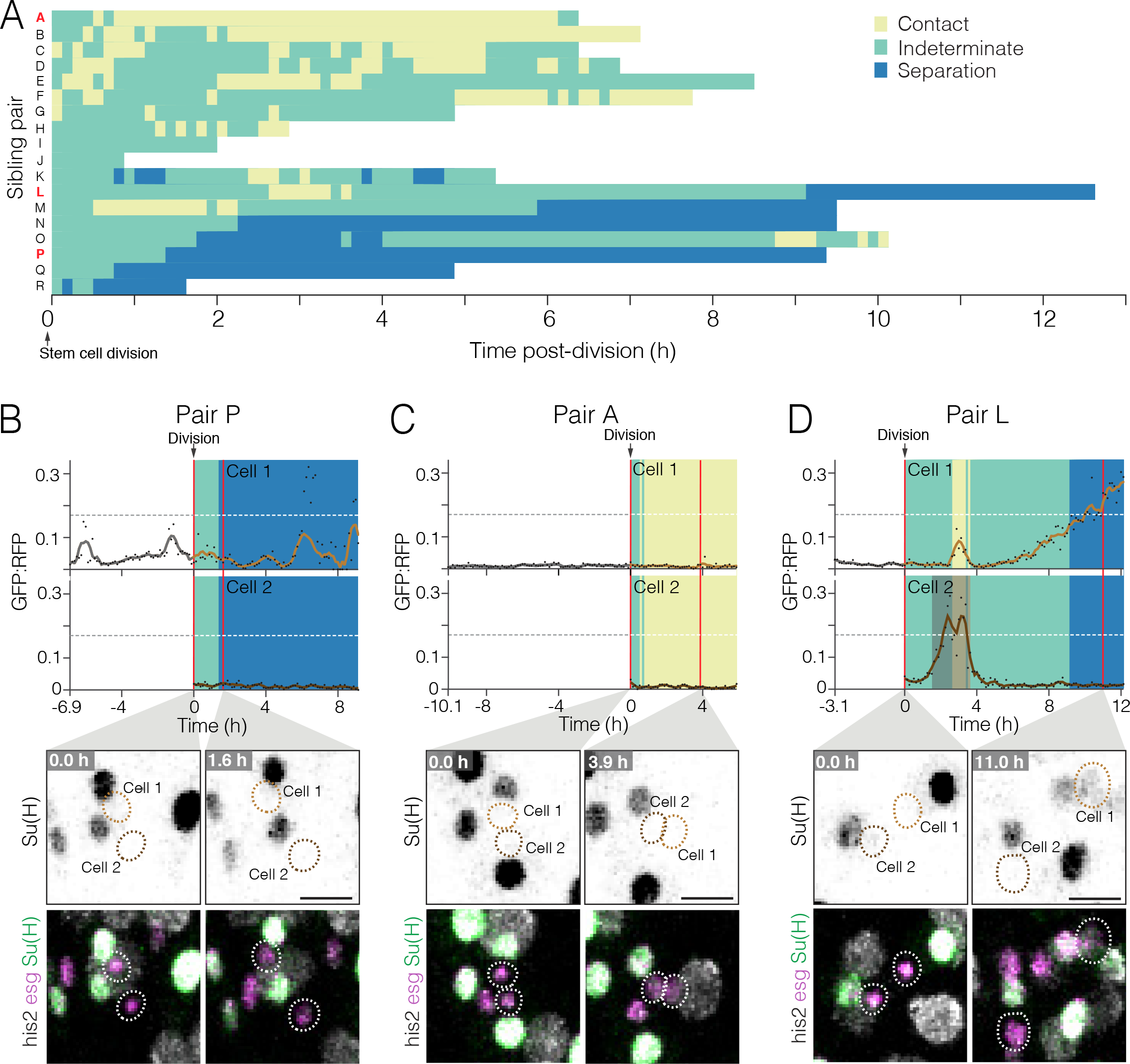
Dynamics of cell contact and Notch reporter activation in sibling cells after birth. **(A)** Contacts between newborn siblings are highly variable. Eighteen pairs of sibling cells (rows A-R) were tracked from birth (t=0.0 h) to the end of imaging. Color shows the likelihood of sibling-sibling contact based on inter-nuclear distance (Fig. S4): Yellow, inferred contact (inter-nuclear distance <6.0 μm); green, indeterminate (inter-nuclear distance =6.0-15.5 μm); blue, inferred separation (inter-nuclear distance >15.5 μm). Pairs are ordered from highest (A) to lowest (P) contact. Pairs A, L, and P (red labels) are featured in Panels C, D, and B, respectively. **(B-D)** Contacts between siblings do not correlate with real-time *GBE-Su(H)-GFP:nls* activation. Graphs show real-time contact status (background colors same as A) and GFP:RFP ratios. Sibling birth is at t=0.0 h. Red vertical lines are the time points shown in the bottom images. **B**, Low-contact pair P does not persistently activate *GBE-Su(H)-GFP:nls*. **C**, High-contact pair A does not persistently activate *GBE-Su(H)-GFP:nls*. **D**, Indeterminate-low contact pair L persistently activates *GBE-Su(H)-GFP:nls* in one sibling. Pair L siblings are in likely contact from t=2.6-3.6 h and are likely separated after t=9.1 h. GFP:RFP of Cell 1 starts to increase at t=4.0 h and reaches the enteroblast threshold at t=10.2 h, a transition period lasting 6.2 h. GFP:RFP of Cell 2 spikes from t=1.5-3.6 h; this spike is not endogenous GFP expression, but rather ‘bleedover’ due to a collision with a mature enteroblast (Video 14). Genotype for all panels: *esgGal4*, *UAS-his2b:CFP*, *Su(H)GBE-GFP:nls*; *ubi-his2av::mRFP*. All scale bars are 10 μm. See Videos 12-14.

Does contact between siblings correlate with Notch activation in real time? To the contrary, both high-and low-contact siblings generally maintained low GFP:RFPs (Fig 6B-C, Videos 12, 13). Of the 36 individual siblings we tracked (Fig. 6A), only one showed persistent activation of Notch (Cell 1 of Pair L, Fig. 6D; Video 14). This particular cell was in likely contact with its sibling for at least one hour, perhaps longer, before its GFP:RFP began to increase (Fig. 6D, t=4.0 h). GFP:RFP climbed to the enteroblast threshold over the subsequent 6.2 h (Fig. 6D, gray background shading), during which time the two siblings likely lost contact (Fig. 6D, t=10.2 h). All other sibling cells remained stem-like, with no persistent Notch activation, until the end of imaging. Had imaging continued, it is possible that additional siblings might have transitioned to enteroblasts. Unfortunately, it is impossible to fathom the influence of sibling contacts on such hypothetical fates. Nonetheless, our present movies imply that contacts between sibling cells after birth do not lead to real-time Notch activation.

### A period of latency after birth?

The temporal kinetics of *GBE-Su(H)-GFP:nls* activation suggest a pattern in which cells delay enteroblast differentiation for multiple hours after birth. In Fig. 6D, 10 hours elapsed between birth and the enteroblast threshold. In Fig. 5D-E, 7-12 hours elapsed between the start of imaging and the enteroblast threshold; furthermore, these four cells were all born before imaging started, so actual times since birth were even longer. Notably, these temporal kinetics could not have been elucidated from fixed samples.

The delay in Notch target activation may explain why we observed only one newborn cell transition to an enteroblast in our movies (Fig. 6D). This frequency was lower than expected based on prior studies of twinspot clones, in which 20-30% of sibling pairs exhibited asymmetric, stem-enteroblast fates (Chen et al., 2015; O’Brien et al., 2011). (Like our movies, these studies examined 2-day midguts, which are undergoing adaptive growth.) The discrepancy between live imaging and clonal analysis could be due to their different time scales. Whereas live imaging tracks cells for hours, twin-spot clones are analyzed after a chase period of days or weeks. Hence, a multi-hour delay in Notch target activation would cause a newborn, incipient enteroblast to be indistinguishable from a stem cell in a movie, but—days later—readily identified as a mature enterocyte in a twin-spot clone.

The root causes of delayed Notch activation can be technical, biological, or both. On the technical side, issues related to sample preparation and imaging cannot be excluded. Temporal factors, such as the time required for membrane activation and nuclear translocation of the Notch receptor (~10 min) and for biosynthesis of GFP (~15 min) (Balleza et al., 2018; Couturier et al., 2012; Housden et al., 2013; Kawahashi and Hayashi, 2010; Vilas-Boas et al., 2011), are unlikely given the multi-hour duration of our movies.

On the biological side, an intriguing possibility is that new cells are refractory to Notch activation for many hours after birth. Mechanisms that impose a period of latency could allow a new-born cell to sample a large variety of signals before ‘choosing’ to either differentiate or self-renew. Each sibling pair will eventually adopt one of three fate combinations: stem-enteroblast, stem-stem, or enteroblast-enteroblast (de Navascués et al., 2012; O’Brien et al., 2011). At the whole-organ level, these three combinations must be collectively balanced to preserve homeostasis (Klein and Simons, 2011). By enabling newborn cells to integrate intrinsic and extrinsic signals, a latent, ‘waiting’ period could ensure that the fates of individual cells are coordinated with the overall needs of the organ.

## CONCLUSION

The *Drosophila* adult midgut is a premier model for organ renewal, but understanding the dynamics of renewal has been hampered by a lack of robust methodology for live imaging. Here, we have presented an imaging platform that captures the midgut in a near-native state within a live animal, yielding movies of exceptional visual quality and duration. In conjunction, we have described a pipeline for comprehensive, 4D movie analysis. We applied this pipeline to our movies for proof-of-principle analyses that corroborated fixed-tissue observations and uncovered new renewal behaviors. These novel findings ranged from time-resolved, single-cell dynamics of division orientation and apical extrusion to large-scale, population-level measurements of Notch activation. The ability to simultaneously span cell- and tissue-level scales over extended imaging periods opens the door to quantitative study of the spatiotemporal complexity of tissue renewal.

In addition to midgut dynamics, our platform offers the opportunity to investigate other biological phenomena in the *Drosophila* abdomen. Animals ingest food during imaging, which enables real-time observation of events, such as colonization by ingested pathogens, that occur in the midgut lumen. By shifting the position of the cuticular window, the platform could also be adapted to study events in other abdominal organs, including ovary, testis, Malpighian tubule, and hindgut. We anticipate that long-term imaging of Drosophila adults will lead to new, dynamic understanding of the cell and tissue behaviors that govern the form and physiology of mature organs.

## Acknowledgements

E.N.S was supported by NSF GRFP DGE-1656518 and Stanford Developmental Biology and Genetics NIH Training Grant 2T32GM00779038. P.M.R. was supported by a Stanford Bio-X Bowes Graduate Fellowship and a Stanford DARE (Diversifying Academia Recruiting Excellence) Fellowship. X.D. was supported by NRSA 1F32GM115065 and a Stanford Dean’s Postdoctoral Fellowship. L.A.K.J. was supported by 1F31GM123736-01and by Stanford Developmental Biology and Genetics NIH Training Grant 2T32GM00779038. This work was supported by NIH R01GM116000-01A1, the Stanford Discovery Fund Innovation Program, and a Center for Biomedical Imaging at Stanford Seed Grant to L.E.O. Confocal microscopy was performed at the Stanford Beckman Cell Sciences Imaging Facility (NIH 1S10OD01058001A1). We thank Benjamin Bolival for the construction of the fate sensor stocks; B. Ohlstein, Y. Inoue, D. Montell, J. de Navascués, and the Bloomington Drosophila Stock Center for fly stocks; T. Larsen and B. Pruitt for assistance with and use of a micro laser cutter; Miriam Goodman for helpful discussions; and C. Cabernard, A. Bardin, J. de Navascués, S. Brantley, and A. Sherlekar for valuable comments on the manuscript.

## Author Contributions

J.M. developed and refined the live imaging platform and analysis pipeline.

J.M., P.M.R. and L.E.O. designed experiments.

J.M. and P.M.R. performed confocal microscopy.

J.M., E.N.S., P.M.R., X.D., and L.E.O. designed quantitative analyses of confocal data.

J.M., E.N.S., P.M.R., L.A.J.K. and S.B. performed quantitative analyses of confocal data.

J.M., E.N.S., L.A.J.K and L.E.O wrote the manuscript.

J.M., E.N.S., P.M.R., S.B., X.D., L.A.J.K and L.E.O. edited the manuscript.

## SUPPLEMENTAL DATA FIGURES S1-4

**Fig. S1.**
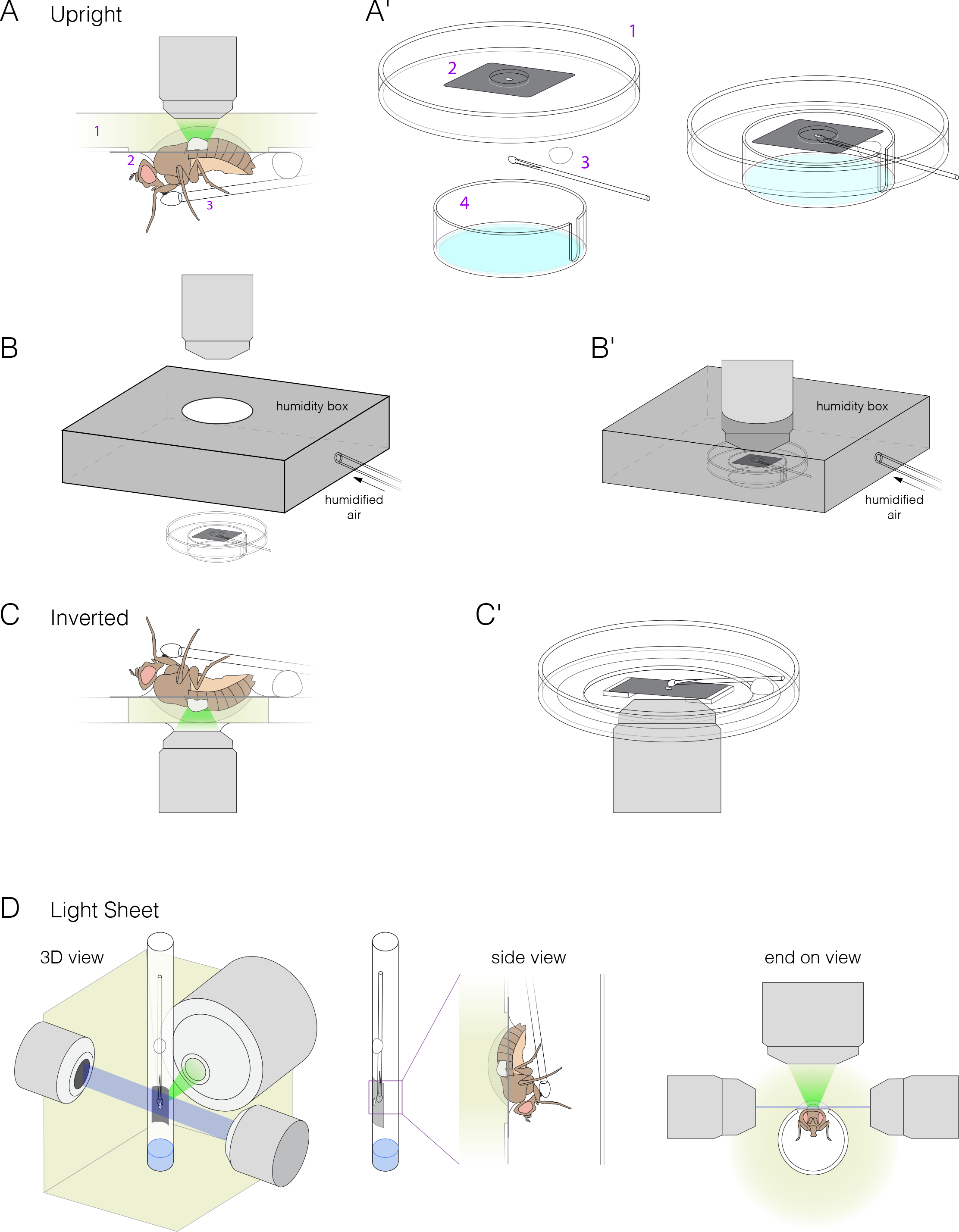
Mounts for upright, inverted, and light sheet microscopes. **(A-B)** Mount for upright microscopes. **A,** Schematic of animal in mount on microscope stage. **A′**, Isometric illustrations of mount components: (1) modified petri dish, (2) metal shim with cutout for *Drosophila* abdomen (Fig. S2), (3) feeder tube, and (4) bottom chamber with wet Kimwipes (light blue). (Bottom chamber is not shown in A.) **B,** Schematic of humidity box that encloses the mount. Unassembled (B) and assembled (B**′**) views are shown. **(C)** Mount for inverted microscopes. The midgut is imaged through a glass-bottom petri dish. To elevate the animal, two spacers are glued to the bottom of the dish, and the metal shim is affixed to the spacers. Media is added to the level of the spacers. **(D)** Mount for light-sheet microscopes. The barrel of a 1-ml syringe is modified to fit the metal shim. The animal and feeder tube are inside the barrel, and the dorsal surface and exposed midgut are outside the barrel. The barrel is submerged in media with one end remaining open to air. 3D, side, and end-on views are shown.

**Fig. S2.**
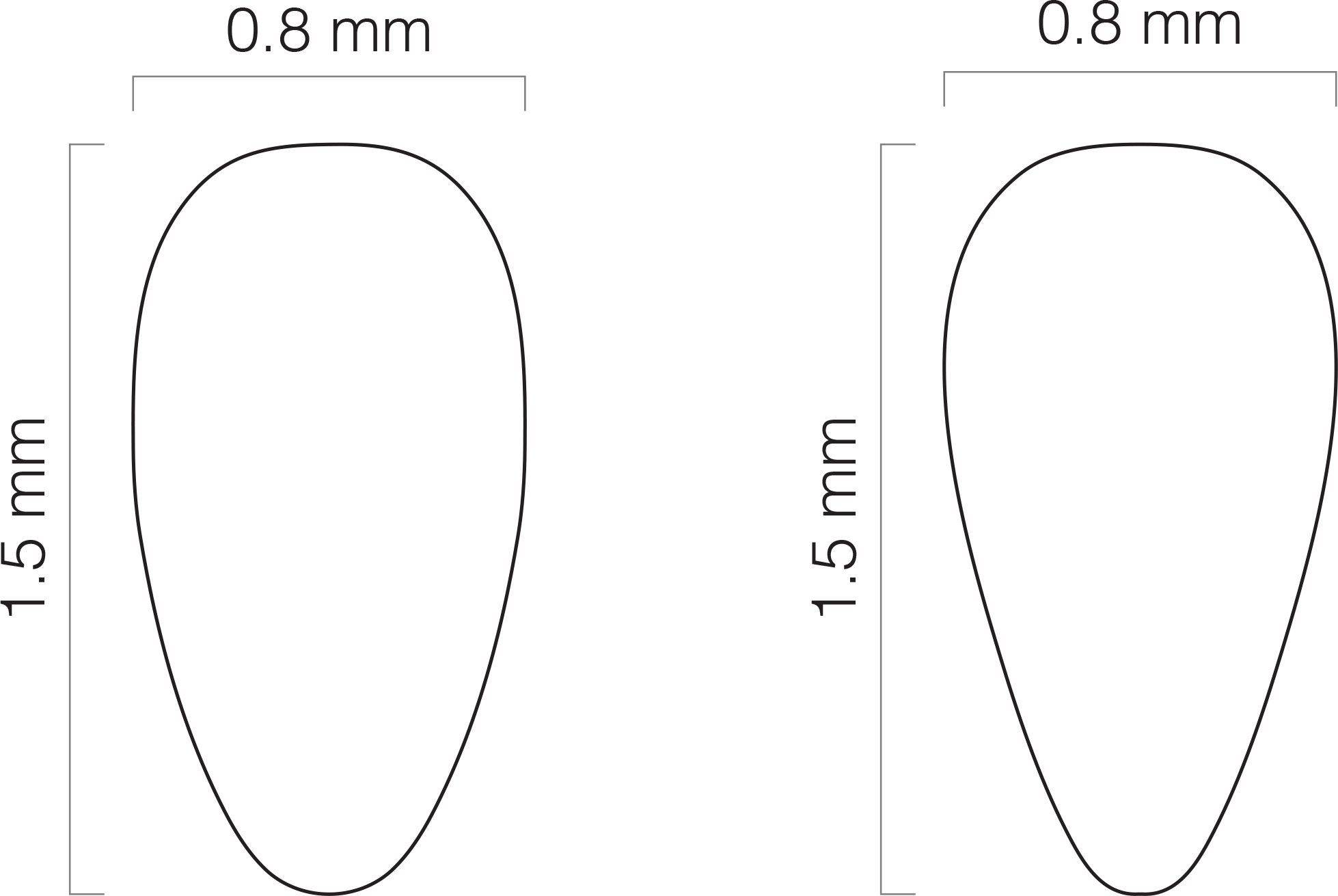
Specifications for abdomen cutouts. The metal shim of the imaging mount includes a cutout through which the dorsal abdomen is inserted. ‘Fat’ (left) and ‘skinny’ (right) cutouts accommodate differently sized female abdomens. This diagram can be used as a CAD file for automated laser cutters.

**Fig. S3.**
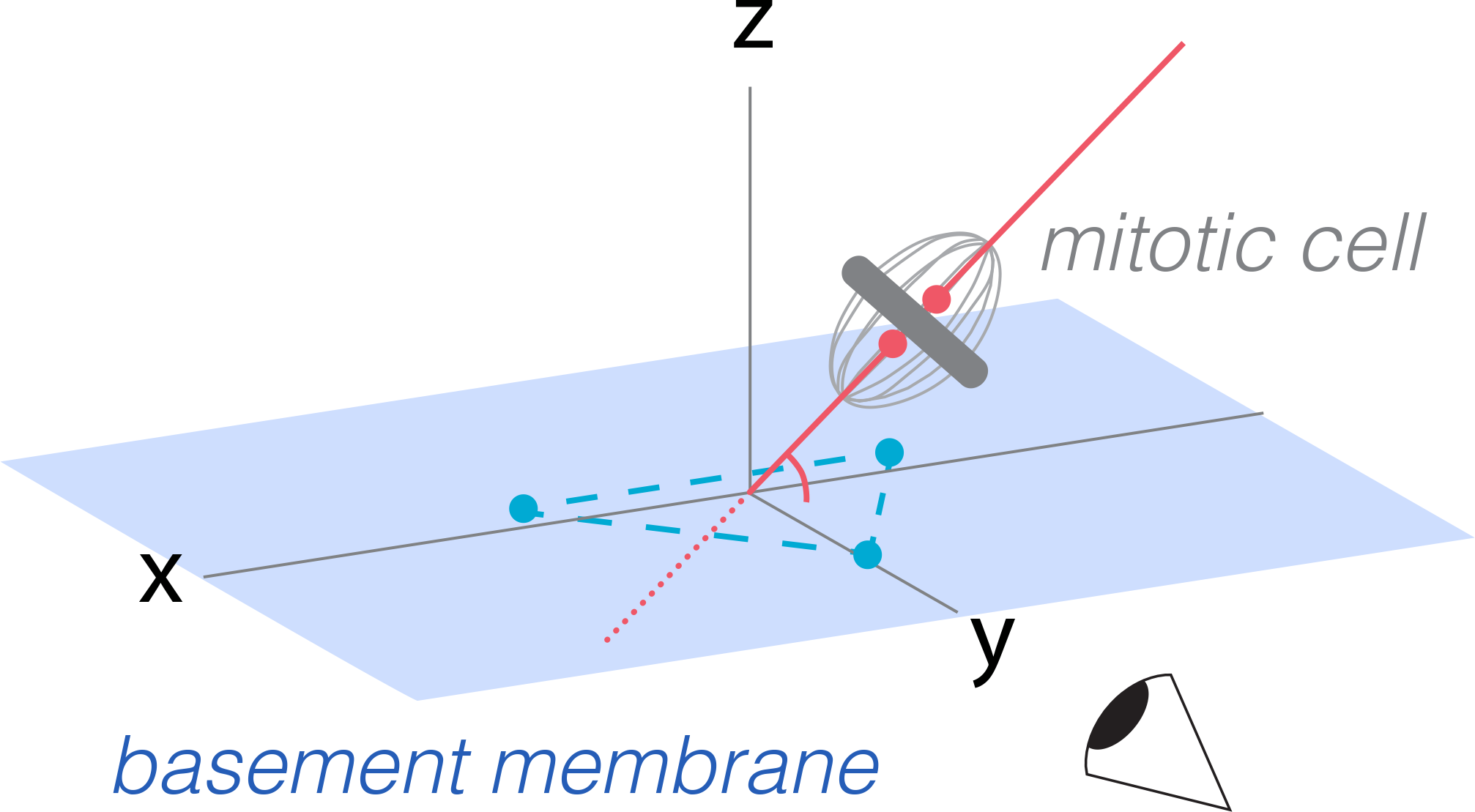
Measurement of horizontal-vertical spindle orientation in space. Horizontal-vertical orientation of the mitotic spindle was measured as the angle at which the presumptive spindle axis intersected a plane tangent to the basal surface of the mitotic cell. Spindle axes and basal planes were determined by examination of volumetric movies in Imaris. To establish coordinates for the spindle axis (red line), two points (red dots) were placed relative to the condensed chromatin (*ubi-his2ab::mRFP*). To establish coordinates for the basal plane (blue rectangle), three points (blue dots) were placed on the basal epithelial surface underlying the spindle. The (x,y,z) coordinates of these five points were input into a vector algebra expression to calculate the horizontal-vertical spindle angle (see Methods).

**Fig. S4.**
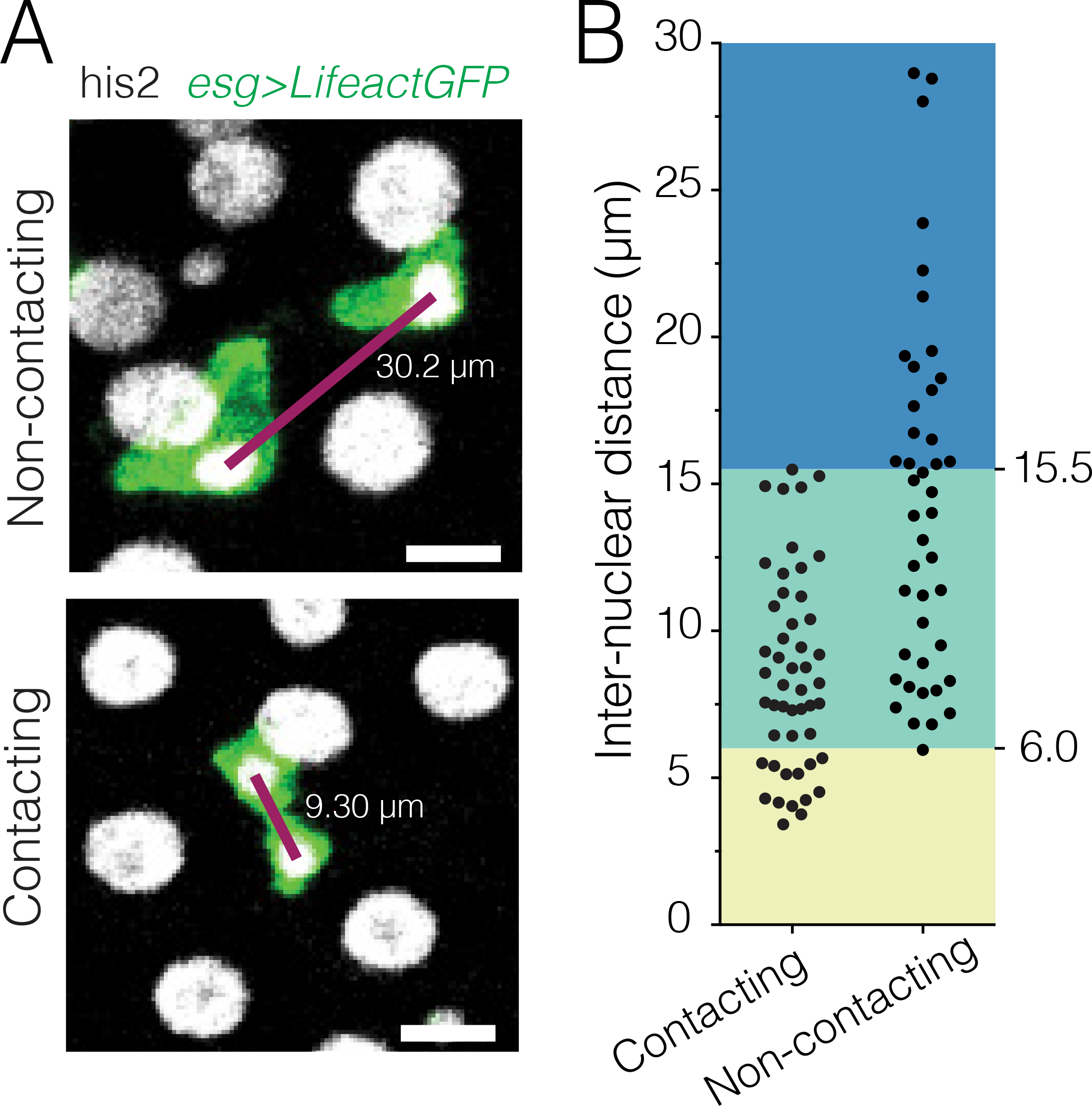
Comparison of cell-cell contact and inter-nuclear distance for live pairs of progenitor cells. **(A)** Examples of contacting and separated progenitor pairs. Contact is revealed using *esg*-driven LifeactGFP (green) to label the actin cytoskeleton of progenitor cells. Inter-nuclear distance (purple lines) is the distance between the centroids of the two nuclei (gray). Images are projections of single time points; however, contact and inter-nuclear distance were assessed in volumetric space. Scale bars, 10 μm. **(B)** Inter-nuclear distances of contacting and separated pairs. All pairs with inter-nuclear distances <6.0 μm (yellow background) are in contact (26% of all contacting pairs). Pairs with inter-nuclear distances from 6.0-15.5 μm (green background) are split between contacting and separated (73% of all contacting pairs; 58% of all separated pairs). All pairs with inter-nuclear distances >15.5 μm (blue background) are separated (42% of all separated pairs). These three ranges are used in Fig. 6 to infer the likely contact behavior of *esg*^+^ sibling pairs that express only nuclear markers (inter-nuclear distance <6.0 μm, likely contact; 6.0-15.5 μm, indeterminate; >15.5 μm, likely separated). n=49 contacting and 43 separated pairs. Pairs were designated as two *esg*^+^ cells that were mutually closer to each other than to any other *esg*^+^ cell and were selected randomly from single time points of four movies. Genotype: *esgGal4*, *UAS-LifeactGFP*; *ubi-his2av::mRFP*.

## CAPTIONS FOR SUPPLEMENTAL VIDEOS 1-14

**Video 1**. Narrated, step-by-step tutorial illustrates an animal being prepared for midgut imaging in the fly mount.

**Video 2**. Fifteen-hour, volumetric movie of the midgut illustrates the wide-field, high-resolution images that are acquired. Numerous physiological contractions of the midgut are evident. An enteric tracheal tube is visible in the lower left. Scale bar, 70 μm.

**Video 3**. After 16 hours of continuous imaging, the animal is alive and responsive.

**Video 4**. Movie clip of midgut before (left) and after (right) stack registration. Before registration, blurred cells from tissue movements are evident during t=20-60 min. After registration, blurring is negligible. Cyan, all nuclei (*ubi-his2ab::mRFP*); yellow, stem cells and enteroblasts (*esg>LifeactGFP*). Each time point is the projection of a confocal z-stack. Scale bar, 20 μm.

**Video 5.** Ten-hour movie of a ‘fate sensor’ midgut (*esgGal4, UAS-his2b::CFP, GBE-Su(H)-GFP:nls; ubi-his2av::mRFP*. See Fig. 2A-B). Nuclei are distinguishable for four midgut cell types: stem cells (red pseudocolor), enteroblasts (yellow-green pseudocolor), enterocytes (gray, polyploid), and enteroendocrine cells (gray, diploid). Each time point is the projection of a confocal z-stack. Scale bar, 20 μm.

**Video 6**. Twelve-hour movie of a single-enterocyte extrusion. The epithelium is oriented with its basal surface toward the microscope objective and its apical surface further away. The basal region of the extruding enterocyte (tan pseudocolor at t=−150, −22.5, 135, 292.5 min) is outlined by a ‘ring’ of E-cadherin::YFP. The ring closes down to a point from t=-105-293 min. The intensity of the ring fluctuates during the first half of closure and becomes consistently bright during the second half. As the ring closes, neighboring cells draw into a rosette. Meanwhile, the nucleus of the extruding cell (yellow pseudocolor) starts to drop apically at t=0 min, hits its deepest luminal position at t=113 min, and recoils from t=113-160 min. Cyan, all nuclei (*ubi-his2av::mRFP*); inverted gray, E-cadherin (*ubi-DE-cadherin::YFP*). Each time point is the projection of a confocal stack. Scale bar, 10 μm.

**Video 7**. Orthoview of same extrusion as Video 6. The nucleus of the extruding enterocyte (magenta) ejects out of the epithelium (t=0-15 min) and penetrates into the lumen (t=15-113 min). It subsequently recoils and eventually comes to rest on the apical epithelium (t=113-293 min). Multicolored line shows the path of nuclear travel over time (violet-yellow color scale; see Fig. 3D for legend). Cyan, all nuclei (*ubi-his2av::mRFP*), gray, E-cadherin (*ubi-DE-cadherin::YFP*). Scale bar, 10 μm.

**Video 8**. Mitosis of a putative stem cell. Green, actin (*esg>LifeactGFP*); yellow, E-cadherin (*ubi-DE-cadherin::YFP*); red, nuclei (*ubi-his2av::mRFP*). Each time point is the partial projection of a confocal stack. Scale bar, 10 μm.

Video 9. Orthoview of a stem cell division with two horizontal-vertical reorientations. The first re-orientation occurs between metaphase (24° at 7.5 min) and anaphase 60° at 15 min). The second re-orientation occurs between anaphase (62° at 22.5 min) and telophase (2° at 30 min). Gray channel, condensed chromatin (*ubi-his2ab::mRFP*). Red line indicates the spindle axis. Cyan line indicates the basal plane, as revealed by the basement membrane stain Concanavalin A-Alexa 647 (not shown). At each time point, the mitotic cell is shown as an orthogonal projection from the vantage of a plane that is parallel to the spindle axis and normal to the basal epithelial plane. For clarity, a clipping plane was applied in the gray channel to exclude an adjacent enterocyte nucleus. Scale bar, 5 μm.

**Video 10**. Division of a stem cell that contacts two enteroblasts. Division orientation aligns with the axis between the two enteroblast nuclei (magenta, *GBE-Su(H)-GFP:nls*). At cytokinesis (t=15-22.5 min), the new daughter nuclei hurl into the enteroblast nuclei, which recoil in response. Gray, stem cell and enteroblast nuclei (*esg>his2b::CFP*). Each time point is the partial projection of a confocal stack. Scale bar, 10μm.

**Video 11**. Real-time enteroblast transition. In the incipient enteroblast (blue dotted circle), *GBE-Su(H)-GFP:nls* is initially undetectable (GFP:RFP=0.014 at t=0.0 h). Over time, its GFP intensity increases, eventually reaching the enteroblast threshold (GFP:RFP=0.18 at t=10.5 h). See Fig. 5D. Right video: Green, *GBE-Su(H)-GFP:nls*.; magenta, stem cell and enteroblast nuclei (*esg>his2b::CFP*); gray, all nuclei (*ubi-his2av::mRFP*). Left video: inverted gray, *GBE-Su(H)-GFP:nls*. Each time point is the partial projection of a confocal stack. Scale bar, 2 μm.

**Video 12**. A low-contact sibling pair (Pair P; Fig. 6A, B) does not activate *GBE-Su(H)-GFP:nls*. Following their birth at t=0.0 h, the two siblings move apart and have likely lost contact by t=1.4 h (in-ter-nuclear distance>15.5 μm; c.f. Fig. S4). The mother stem cell is indicated by the blue dotted circle at t=-1.0 h; the two siblings are indicated by the two blue dotted circles at t=0.0 and t=9.2 h. No GFP expression is apparent in either sibling. Right video: Green, *GBE-Su(H)-GFP:nls*; magenta, stem cell and enteroblast nuclei (*esg>his2b::CFP*); gray, all nuclei (*ubi-his2av::mRFP*). Left video: inverted gray, *GBE-Su(H)-GFP:nls*. Each time point is the partial projection of a confocal stack. Scale bar, 5 μm.

**Video 13**. A high-contact sibling pair (Pair A, Fig. 6A, C) does not activate *GBE-Su(H)-GFP:nls*. Following their birth at t=0.0 h, the two siblings remain in likely contact (inter-nuclear distance <6.0 μm; c.f. Fig. S4) for at least 6.0 h. The mother stem cell is indicated by the blue dotted circle at t=-1.2 h; the two siblings are indicated by the two blue dotted circles at t=0.0 and t=6.0 h. No GFP expression is apparent in either sibling. Right video: Green, *GBE-Su(H)-GFP:nls*; magenta, stem cell and enteroblast nuclei (*esg>his2b::CFP*); gray, all nuclei (*ubi-his2av::mRFP*). Left video: inverted gray, *GBE-Su(H)-GFP:nls*. Each time point is the partial projection of a confocal stack. Scale bar, 5 μm.

**Video 14**. A sibling pair exhibits asymmetric Notch activation (Pair L, Fig. 6A, D). Following their birth at t=0.0 h, the two siblings are in likely contact from t=2.6-3.6 h, are in indeterminate contact from t=3.6-9.0 h, and are likely separated after t=9.0 h. The mother stem cell is indicated by the blue dotted circle at t=-1.0 h. The two siblings are indicated by the two blue dotted circles at t=0.0 and 12.2 h. A single blue dotted circle at t=10.2 h indicates when the Notch-activated sibling crosses the enteroblast threshold (GFP:RFP=0.17; c.f. Fig. 6D). This sibling exhibits nascent GFP signal at 4.0 h and increases in GFP intensity for the rest of the movie. The other sibling does not exhibit detectable GFP, but from t=1.5-3.6 h, it collides with a high-GFP enteroblast (orange dotted circle), which causes GFP signal to ‘bleed over’ in the GFP:RFP analysis (Fig. 6D). Right video: Green, *GBE-Su(H)-GFP:nls*; magenta, stem cell and enteroblast nuclei (*esg>his2b::CFP*); gray, all nuclei (*ubi-his2av::mRFP*). Left video: inverted gray, *GBE-Su(H)-GFP:nls*. Each time point is the partial projection of a confocal stack. Scale bar, 5 μm.

## MATERIALS AND METHODS

### *Drosophila* husbandry

Fly stocks obtained from other sources:

*esgGAL4* (Kyoto DGRC)
*ubi-his2av::mRFP* (BL23650)
*UAS-LifeactGFP* (BL35544)
*UAS-his2b::CFP* (Yoshihiro Inoue) (Miyauchi et al., 2013)
*GBE-Su(H)-GFP:nls* (Joaquin de Navascués) (de Navascués et al., 2012)
*act5c-spaghetti squash::GFP* (Denise Montell)
*ubi-E-cadherin::YFP* (Denise Montell) (Cai et al., 2014)

Generated stocks:

*act5c-spaghetti squash::GFP; ubi-his2av::mRFP*
*esgGal4, UAS-his2b::CFP, GBE-Su(H)-GFP:nls; ubi-his2av::mRFP*
*esgGAL4, UAS-his2b::CFP, GBE-Su(H)-GFP:nls/CyO; ubi-his2av::mRFP*
*esgGal4, UAS-LifeactGFP; ubi-his2av::mRFP*
*esgGAL4, GBE-Su(H)-GFP:nls; UAS-his2ab::mRFP*

Adult female Drosophila 2.5 days post-eclosion were used in all movies except Video4old, which used a female 7 days post-eclosion. Females were collected 0-4 h post-eclosion, placed in vials with males, and maintained at 25 °C. Flies were fed a diet of standard cornmeal molasses food supplemented with yeast paste (1 g/1.4 ml H_2_O).

### Fly mounts for extended live imaging on upright, inverted, and light-sheet microscopes

We designed three types of fly mounts that enable dorsal exposure of the midgut while stabilizing the live intact animal. For upright and inverted microscopes, our design is a modification of a previously published mount for imaging of adult *Drosophila* brains (Seelig et al., 2010).

### Upright mount

To prepare the upright mount (Fig. S1A-B), a stainless steel shim of 0.001 in thickness (Trinity Brand Industries, 612H-1) was cut into 19 x 13mm rectangles. From these rectangles, abdomen-sized cutouts were excised either by hand using an 18-gauge PrecisionGlide needle (Becton Dickinson, 305196), or by laser cutting using a micro laser cutting system (see Fig. S2 for CAD file). In addition, we prepared 60-mm petri dishes (Fisher, FB0875713A) with a hole 10 mm in diameter drilled into the bottom. Each shim was glued onto the base of a 60-mm petri dish with clear silicone glue (DAP, 00688) and allowed to dry overnight.

The mount includes a feeder tube to provide the animal with liquid nutrients during imaging. We found that the feeder tube was essential for prolonged survival of the animal. Feeder tubes were made from 20-μl capillary tubes (Sigma-Aldrich, P0799), which were cut into 38-mm sections. Using a tungsten wire with a small hook bent at the end, a bit of cotton (Fisher Scientific, 22-456-880) was pulled through one end of the capillary tubing to form a feeder wick. Attachment of the feeder tube to the mount is described below.

A protective bottom chamber (Fig. S1A′) enclosed the ventral side of the animal during imaging to prevent desiccation. To create the chamber, a 3-mm wide channel was drilled down the wall of a 35-mm petri dish (Olympus Plastics, 32-103). Kimwipes (4-ply rounds, Fisher Scientific, 06-666) were cut and placed in the bottom of the humidity chamber to be soaked with water before use.

### Inverted mount

The inverted mount (Fig. S1C-D) was similar to the upright mount, but used a glass bottom petri dish with two 1-mm spacers glued approximately 10 mm apart. Spacers were made from cut pieces of glass microscope slides (63720-05, Electron Microscopy Sciences) and were adhered with silicone glue. The same metal shim was used as with the upright mount, but was not affixed to the dish until the animal was glued and its gut stabilized (see below). Once the animal was prepared, the mounting shim was positioned with animal’s dorsal side toward the glass bottom of the dish and glued to the spacers using KWIK-SIL adhesive silicone glue (World Precision Instruments, 60002).

### Light-sheet mount

Zeiss light sheet systems require a submersible chamber. To create such a chamber, we used the barrel of a 1-ml syringe in which one end was open to air (Fig. S1D). A 5-mm x 8-mm square was cut into the side of the syringe barrel, and a metal shim with abdominal cutout was affixed to the square window using silicone glue. A second window was cut into the opposite side of the barrel to provide physical access for mounting the animal and feeder tube inside. Once the animal and feeder tube were in place, the access window was sealed using a second metal shim and KWIK-SIL glue (World Precision Instruments, 60002). The bottom end of the syringe was sealed with dental wax (Surgident, 50092189) and the barrel was submerged in media in the Zeiss sample chamber. In this manner, the midgut was bathed in media during imaging while the animal’s head and ventral surface remained in an open-air environment.

## Composition of imaging media and agarose

Media for midgut imaging was based on prior recipes for *Drosophila* organ culture *ex vivo* (Morris and Spradling, 2011; Zartman et al., 2013). Schneider’s Insect Media (Sigma-Aldrich, S0146) was supplemented with 5% FBS (Sigma-Aldrich, F4135), 5% fly extract (DGRC) (Currie et al., 1988), 100 μg/mL human insulin (Sigma-Aldrich, I0516) and 0.5% penicillin-streptomycin (Invitrogen, 15140). (Without antibiotics, the imaging media became visibly contaminated after several hours of imaging.) Insulin was added fresh on the day of imaging.

Low-melting point agarose was used to stabilize the midgut during imaging. To make the agarose, 2-Hydroxyethylagarose (Sigma-Aldrich, A4018) was mixed with Schneider’s to make a 6% w/v slurry. The slurry was heated to 65 °C to melt the agarose, mixed thoroughly and separated into 25μL aliquots that were stored at 4 °C. The day of imaging, aliquots were heated to 65 °C, mixed 1:1 with a 2x concentration of imaging media, and applied to midguts as described below.

## Animal preparation

A narrated tutorial video (Video 1) provides step-by-step instructions for mounting the animal and exposing the midgut. Wings were broken off near the hinge using forceps. Flies were placed in a microfuge tube on ice for at least 1 h before being glued dorsal side down to the fly mount (Fig. S1) using KWIK-SIL glue. To optimize access to region R4 of the gut, the fly was tilted toward its left side when glued to the mount. For long-term survival of the animal, its right-side spiracles were kept open (Video 1). After the glue had dried, the feeder tube, filled with 10% sucrose (w/v) in H_2_O, was secured to the fly mount with dental wax (Surgident, 50092189) and positioned such that the cotton wick was in reach of the animal’s proboscis. The protective bottom chamber was attached to the bottom of the petri dish with masking tape (Fig. S1A′).

To expose the midgut, a window was cut into the dorsal cuticle as follows: First, a drop of imaging media was placed onto the dorsal cuticle. Next, portions of cuticular segments A1 and A2 were excised using Dumont #55 forceps. In the majority of animals, this excision exposed the looped midgut region R4a-b/P1-2 (Buchon et al., 2013; Marianes and Spradling, 2013). The loop was gently coaxed using forceps to protrude slightly out of the window. The imaging media was temporarily removed, and a drop of the agarose mixture (described above) was applied to the exposed loop and allowed to solidify. Once the agarose had hardened, a drop of media was added on top of the agarose to avoid desiccation. Between steps, the setup was placed on ice to minimize animal movement. The bottom of a 60-mm petri dish was used to cover the mounted animal until ready for placement on the microscope.

In some experiments, Concanavalin A-Alexa647 (Invitrogen, C21421) was used to stain the basement membrane. A stock solution of 5 mg/mL Concanavalin A in 0.1 M sodium bicarbonate was diluted 1:200 in 1x imaging media to obtain a final working concentration of 25 μg/mL. A drop of this media was placed on the dorsal cuticle prior to cutting the cuticle window. Agarose and subsequently added media did not contain Concanavalin A.

## Microscopy

An upright Leica SP5 multi-photon confocal microscope and a 20x water immersion objective (Leica HCX APO L 20x NA 1.0) were used to acquire the movies that were analyzed in this study. The microscope was controlled via a Leica CTR6500 controller card on a Z420 (Hewlett Packard) workstation with 16GB memory and a Xeon CPU E5-1620 (Intel) running Windows 7 Pro and Leica Application Suite: Advanced Fluorescence (LAS AF, v.2.7.3.9723). In addition, an inverted Leica SP8 confocal microscope with a 20x oil immersion objective (Leica HC PL APO IMM CORR CS2 NA 0.75) and a Zeiss light sheet Z.1 with a 20x water immersion objective (Zeiss light sheet detection optics 20X NA 1.0) were used to test the fly mounts for these microscope setups. For upright and inverted setups, a humidity box was assembled around the lens and the specimen to prevent desiccation (Fig. S1B). The humidity box was formed from a pipette box lid with a hole for the lens and an inlet tube for humidified air. The box was connected to a 500-ml Pecon humidification bottle containing distilled water, and humidified, ambient air was piped into the box via a Pecon CTI-Controller 3700. In addition, for upright setups, the Kimwipes in the protective bottom chamber were saturated with distilled water. For upright setups, 2-3 ml of imaging media were added to the sample, spanning the distance between the exposed midgut and the water immersion objective. Movies were captured at room temperature (20-23 °C). Confocal stacks were acquired with a z-step of 2.98 μm and typically contained ~35 slices.

## Movie registration and cell masking in Fiji

After acquisition, movies were processed on a Mac Pro computer (OS X 10.8.5) with a 3.2 GHz quad-core Intel Xeon processor and 20GB memory. LIF files (*.lif) from LAS AF were uploaded into Fiji as a hyperstack for registration. To correct for X-Y drift, movies were converted to RGB files and processed with the Fiji plugin StackReg (Arganda-Carreras et al., 2006). To correct for global volume movements, movies were processed with the Fiji plugin Correct 3D Drift (Parslow et al., 2014).

For identification of individual cells, intensity thresholding of the ubiquitously expressed nuclear marker *ubi-his2av::RFP* was used to segment out individual cell nuclei. In fate sensor movies (c.f. Fig. 2), intensity thresholding for CFP (stem cells and enteroblasts; *esg>his2b::CFP*) and GFP (enteroblasts; *GBE-Su(H)-GFP:nls*) was applied to define masks of nuclei within each channel. Using Fiji’s image calculator function, these masks were used to isolate the His2av::RFP-marked nuclei for individual cell types. Specifically, after registration (Fig. 1E), channel masks were generated in Image J to digitally isolate stem cells (CFP channel minus GFP channel), enteroblasts (GFP channel), and mature enterocytes and enteroendocrine cells (RFP channel minus CFP channel). To digitally isolate enterocytes and enteroendocrine cells, whose populations were defined by the absence of *esg>his2av::CFP*, the His2b::CFP-marked progenitor nuclei were eliminated using the subtraction function in Fiji’s image calculator.

Masked nuclei for these three populations were added to the raw hyperstack as unique channels for use in Bitplane Imaris (see below). The two types of mature cells, enterocytes and enteroendocrine cells, were then distinguished by a size filter in Imaris; nuclei >113 um^3^ were classified as enteroendocrine, whereas nuclei >113 um^3^ were classified as enterocyte. To maintain metadata structure for 4D Imaris analysis (see below), final movies were exported to OME-TIFF format (*.ome.tif) using the BioFormats plugin in Fiji.

## Cell identification and tracking in Imaris

To perform cell tracking, Fiji processed stacks of midgut movies were opened in Bitplane Imaris from OME-TIFF format (*.ome.tif) files. Once converted to an Imaris *.ims file, the 4D volumes were visually inspected using the Surpass module to verify the accuracy of image processing and file conversion. Cell segmentation was then performed by applying the Surface Recognition Wizard module to the masked cell channels generated in Fiji (see above). Final products were visually compared to raw channels to confirm cell-type recognition.

Cell surfaces were tracked using the Brownian motion tracking algorithm. We found that automatic tracking accurately identified ~75% of cells. Visual inspection was used to correct errors. Once cell recognition was verified for all cells and time points, individual cell statistics were exported as either a Microsoft Excel file or a comma-separated-value file. The data were then imported into Mathematica or MATLAB for quantitative analysis.

For Figures 5 and 6 and their associated data, a modification of the above protocol was used. To identify cells that transitioned over time from a stem-like state (GFP:RFP≤0.17) to an enteroblast state (GFP:RFP>0.17), cells expressing *esg>his2b::CFP* were identified in Imaris. Their GBE-Su(H)-GFP:nls intensities and nuclear volume were determined at each time point. Cells exhibiting increasing GFP intensities were identified and selected for further analysis.

## Spatiotemporal analyses of enterocyte extrusion

***Analyses of E-cadherin::YFP ring***: Extruding enterocytes were identified by visual inspection. To measure dynamics of the E-cadherin::YFP ring (Fig. 3B-E), Fiji-processed movies were opened from OME-TIFF files (*.ome.tif) in Bitplane Imaris and viewed as 3D volumes using the Surpass module. Vertices of the E-cadherin::YFP ring that outlined the extruding cell were identified by visual inspection at each movie time point. The Measurement Points tool was used to place a polygon-mode measurement point at each vertex. In addition, a plane representing the basal epithelium was defined by selecting three Measurement Points on the basal epithelial surface underlying the extruding cell. To identify the position of the basal surface, we used either the basement membrane stain Conconavalin-A-Alexa647 or the background fluorescence of enterocyte cytoplasms when movies were digitally overexposed. The spatial coordinates of all these measurement points were exported as comma-separated values and imported into MATLAB.

To map the ‘footprint’ of the ring in the epithelial plane (Fig. 3B), the coordinates of the ring vertices were connected with a line, and the resulting polygon was projected onto the basal plane for each time point. The polygon was color-coded according to its time point in the movie.

To calculate the cross-sectional area of the ring (Fig. 3C), the centroid of the polygon was triangulated using its vertices. The area of each component triangle was calculated as half of the cross product of the two vectors formed by the centroid and the two adjacent vertices. The area of the ring was calculated as the sum of the areas of each component triangle. Ring areas were calculated for each movie time point.

To determine the apical-basal position of the E-cadherin::YFP ring, we calculated the orthogonal distance from the centroid of the ring to the basal plane. This distance was calculated as the dot product of two vectors: the unit normal vector of one of the basal measurement points, and a vector from the centroid to basal measurement point that was used as the origin of the unit normal vector.

***Analyses of the extruding nucleus***: We defined the duration of nuclear extrusion (Fig. 3F) as the length of time that the extruding cell’s nucleus was moving apically. To determine this duration, Fiji-processed movies were opened from OME-TIFF files (*.ome.tif) in Bitplane Imaris and viewed as 3D volumes using the Surpass module. Nuclei of extruded enterocytes were digitally isolated via clipping planes and viewed in cross section. Nuclei of enterocytes surrounding the extruding cell were used to establish the baseline level of the epithelium. The duration of nuclear extrusion was measured from the time point for which apical movement of the nucleus was first apparent to the time point of maximal apical displacement from the baseline. The centroid of the nucleus was calculated from surface-recognized objects in Imaris. The distance of the nucleus from the basal surface (Fig. 3E) was calculated as the orthogonal distance from the centroid of the nucleus to the basal plane, as defined by the basal epithelium reference points described above.

## Spatiotemporal analyses of stem cell mitoses

***Mitotic duration***: Mitoses were identified by visual inspection of maximum intensity z-projections in Fiji and confirmed in Bitplane Imaris using the Surpass module for 3D visualization. To calculate the durations of individual mitoses (Fig. 3H), we designated the start point as the initiation of nuclear condensation in the mother cell and the end point as the decondensation of the two sets of daughter chromosomes.

***Mitotic index***: To calculate mitotic index, we used 9 movies of *ubi-his2av::mRFP*-expressing midguts that each contained at least one identifiable division. Movies were processed in Fiji, and nuclei were identified and tracked in Imaris as described above. Mitotic index was calculated as TM/TSC, where TM is the sum of the durations of 39 individual mitoses in the 11 movies, and TSC is the sum of the durations of ‘screen time’ for all the stem cells in the same movies. Stem cell ‘screen time’ was calculated as the product of the number of stem cells at t=0 in a particular movie and the duration of that movie. (On occasion, stem cells disappeared or appeared over the course of a movie; however, these events were infrequent and are not included in our calculations.) To determine the number of stem cells in a movie at t=0, we used one of two approaches. For midguts that expressed *esg>his2av::CFP* and *GBE-Su(H)-GFP:nls* in addition to *ubi-his2av::mRFP* (2 of 9 movies), stem cells were identified as CFP+ cells with GFP:RFP<0.17 (c.f. Fig. 5), and the number of stem cells was counted following Imaris surface recognition as described above. For midguts in which stem cells were not identifiable through specific markers (7 of 9 movies), the number of stem cells was estimated as 20% of total, *ubi-his2av::mRFP*-expressing cells (de Navascués et al., 2012; O’Brien et al., 2011).

***Horizontal-vertical orientation***: To measure the real-time horizontal-vertical orientations of dividing cells (Fig. 4A-E). Fiji-processed movies were opened as OME-TIFF files (*.ome.tif) in Bit-plane Imaris. For each mitotic cell, the positions of the spindle poles and of the basal epithelial surface were determined at each time point between the start and end of mitosis (Fig. S3). Spindle pole positions were inferred from the morphology of the condensed chromosomes and marked using the Measurement Points tool. A plane representing the basal epithelium was defined using the Measurement Points tool to place three points on the basal epithelial surface underlying the spindle. To identify the position of the basal surface, we used either the basement membrane stain Con-conavalin-A-Alexa647 or the background fluorescent signal of enterocyte cytoplasms made visible when movies were digitally overexposed.

The coordinates of spindle poles and basal planes in 3D space were exported as Excel files and opened in Mathematica. To calculate the spindle angle, we defined two vectors: the ‘spindle pole vector’, which was the difference between the coordinates of the two spindle poles, and the ‘basal plane vector’, which was the cross product of two vectors determined from the three points defining the basal plane. The spindle angle was calculated as the dot product of the spindle pole vector and the basal plane vector.

***Longitudinal-circumferential orientation***: To measure longitudinal-circumferential orientation of dividing cells (Fig. 4F-H), movies were analyzed as maximum-intensity projections in Fiji. Longitudinal and circumferential axes were determined for each mitotic cell by visual inspection, based on the local morphology of the midgut tube and the orientation of the ellipsoid nuclei of surrounding enterocytes. Division orientation was measured at the time point when we observed de-condensation of the daughter chromosomes, an event signifying the end of mitosis. To calculate longitudinal-horizontal orientation, we used the Fiji Angle Tool, which measures an angle defined by two vectors formed from three points. One vector was defined by the difference between the positions of the two daughter cells, and the other vector was defined by the longitudinal axis of the midgut tube.

***Orientation relative to neighboring enteroblasts***: We identified mitoses in which the dividing cell contacted either one enteroblast or two enteroblasts using visual inspection. To determine the spatial coordinates of the mitotic cells and the enteroblasts, Fiji-processed movies were opened as OME-TIFF files (*.ome.tif) in Bitplane Imaris. Nuclei were recognized using the Surface Recognition Wizard. The 3D coordinates of the relevant cells were exported into MATLAB.

Division orientation relative to neighboring enteroblasts (Fig. 4I-K) were calculated as follows: For mitotic cells contacting one enteroblast, we designated the daughter cell closer to the enteroblast as ‘D1’ and the other daughter cell as ‘D2’. We defined an ‘enteroblast-D1 vector’ as the difference between the coordinates of the enteroblast and D1. We defined a ‘D1-D2 vector’ as the difference between the coordinates of D1 and D2. To calculate the division angle, we computed the dot product of the enteroblast-D1 vector and the D1-D2 vector.

For mitotic cells contacting two enteroblasts, we designated the reference enteroblast as the enteroblast whose nucleus was closer to the mother stem cell nucleus prior to division. D1 and D2 daughters were determined relative to that enteroblast following the procedure detailed above.

Mitotic cells that contacted enteroblasts were excluded from analyses of horizontal-vertical and planar orientations.

## Quantitative assessment of Notch reporter activation

Activation of the Notch reporter *GBE-Su(H)-GFP:nls* was measured in movies of fate sensor midguts (*esgGal4, UAS-his2b::CFP, GBE-Su(H)-GFP:nls; ubi-his2av::mRFP*) (Figs. 5 and 6). As described above, Fiji-processed movies were opened as OME-TIFF files (*.ome.tif) in Bitplane Imaris, and surface recognition was performed to identify individual cell nuclei using the Add New Surfaces function in the Surpass Module. To quantify *GBE-Su(H)-GFP:nls* activation, we calculated the normalized ratio of GFP:nls and His2av::mRFP intensities as follows: (1) To generate normalized intensity values, raw intensity values for GFP and RFP of single cells at each individual time point were determined from cell nuclei. These raw intensities were exported to MATLAB and divided by the maximum intensity in that movie to yield normalized intensities. (2) The ratio of normalized GFP:RFP intensities was calculated for each cell at each time point. The resulting real-time, normalized GFP:RFPs enabled quantitative comparison of *GBE-Su(H)-GFP:nls* expression between different cells, at different times, or across different movies.

## Calculating inter-nuclear distances of progenitor pairs

To perform the initial analysis of inter-nuclear distances for contacting and non-contacting progenitor pairs (Fig. S4), we used movies of midguts with genotype *esgGal4, UAS-LifeactGFP; ubi-His2av::RFP*. For this analysis, two *esg*^+^ cells were designated as a pair if they were mutually closer to each other than to any other *esg*^+^ cell. Pairs were selected randomly from single time points in four separate movies. Movies were examined in 4D using the Surpass Module in Imaris, and *esg*^+^ pairs were identified as either contacting or non-contacting based on their LifeactGFP signal. To determine the inter-nuclear distance of a pair, the (x,y,z) coordinates for centroid of each nucleus was determined based on surface recognition in Imaris. The distance D between the two centroids was calculated using the equation 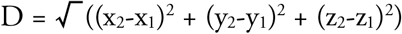.

To determine the inter-nuclear distances for sibling pairs with known birth times (Fig. 6), we used movies of ‘fate sensor’ midguts (*esgGal4, UAS-his2b::CFP, GBE-Su(H)-GFP:nls; ubi-his2av::mRFP*). Following a stem cell division, the inter-nuclear distance of the two siblings at each movie time point was calculated as described above.

